# Optimizing mechanostable anchor points of engineered lipocalin in complex with CTLA-4

**DOI:** 10.1101/2021.03.09.434559

**Authors:** Zhaowei Liu, Rodrigo A. Moreira, Ana Dujmović, Haipei Liu, Byeongseon Yang, Adolfo B. Poma, Michael A. Nash

## Abstract

We used single-molecule AFM force spectroscopy (AFM-SMFS) to screen residues along the backbone of a non-antibody protein binding scaffold (lipocalin/anticalin), and determine the optimal anchor point that maximizes binding strength of the interaction with its target (CTLA-4). By incorporating non-canonical amino acids into anticalin, and using click chemistry to attach an Fgβ peptide at internal sequence positions, we were able to mechanically dissociate anticalin from CTLA-4 by pulling from eight different anchoring residues using an AFM cantilever tip. We found that pulling on the anticalin from residue 60 or 87 resulted in significantly higher rupture forces and a decrease in *k*_*off*_ by 2-3 orders of magnitude over a force range of 50-200 pN. Five of the six internal pulling points tested were significantly more stable than N- or C-terminal anchor points, rupturing at up to 250 pN at loading rates of 0.1-10 nN sec^-1^. Anisotropic network modelling and molecular dynamics simulations using the Gō-MARTINI approach explained the mechanism underlying the geometric dependency of mechanostability. These results suggest that optimization of attachment residue position for therapeutic and diagnostic cargo can provide large improvements in binding strength, allowing affinity maturation without requiring genetic mutation of binding interface residues.

## Introduction

Mechanical anisotropy refers to the variety of mechanical responses that manifest when force is applied to macromolecules from different directions. A classic example is the mechanically induced dissociation of double stranded DNA/RNA double helices and hairpins^1–3^. Depending on the pulling points, DNA can be unzipped at low forces (∼30 pN) where the base paired hydrogen bonds are broken in series, or sheared apart at high forces (>50 pN) where the hydrogen bonds are broken in parallel. Since the sequence is identical in both scenarios, the hybridization energy is equal and the effect is entirely attributable to an anisotropic mechanical response of the double helix. This mechanical property of DNA has been applied to build force hierarchies, enabling one-by-one assembly of complex molecular patterns on surfaces using single-molecule cut-and-paste with the atomic force microscope^4^.

Folded protein domains similarly exhibit a heterogeneity of mechanical responses depending on the pulling geometry. This was shown for several folded domains including GFP^5^, ubiquitin^6^, E2lip3^7^ and GB1^8^. A recent study also reported application of force from an internal sequence position to transmembrane bacteriorhodopsin to dislodge transmembrane helices in a defined order from the membrane and understand intermediate folding states in that system^9^. The force required to dissociate receptor-ligand binding interfaces has also been shown by our group and others to strongly depend on whether the receptor is pulled from the N- or C-terminus, as was shown for cohesin-dockerin and streptavidin-biotin systems^10–12^. Differences in shear *vs*. zip geometry have also been reported for protein-based coiled coils^13^. For protein-protein interactions, however, to the best of our knowledge all previous studies were limited to comparing N- and C-terminal anchor points.

Structure-based heuristics are already in place for predicting the mechanical stability of folded domains stretched between their N- and C-termini^14–17^. In general, high alpha helix content is associated with low unfolding forces and long unfolding barrier positions, while high beta strand content is associated with protein folds that are generally more mechanically robust with short unfolding barrier positions. Conserved structural motifs referred to as mechanical clamps are also known to impart mechanostability to folded domains^18–20^. However, these general trends may change when alternative anchor points are considered. Furthermore, for protein-protein interactions no such heuristics are available. It remains unclear how pulling geometry modulates the binding strength of protein-protein complexes. Single-molecule AFM force spectroscopy (AFM-SMFS) provides a powerful tool to study the force response of protein-protein complexes. State-of-the-art AFM setups are able to precisely control the pulling geometry of receptor-ligand systems and measure a wide range of forces^21–23^.

For applications in targeted drug delivery using nanoparticles, liposome or engineered viruses, the mechanical force that a receptor molecule can withstand while remaining bound to its target ligand is a potentially valuable optimization parameter. One current trend in biotherapeutics is exploring beyond full length IgG antibodies and into the realm of non-antibody scaffolds. Compared to conventional monoclonal antibodies, non-antibody scaffolds are smaller in size, easier to manufacture and exhibit low immunogenicity^24,25^. Scaffolds such as anticalins, DARPins and adnectins can be diversified at the genetic level and evolved in vitro to bind many different molecular targets^26^. Anticalins are a class of non-antibody scaffolds derived from naturally occurring lipocalins^27^. The anticalins share homologous backbone structures (**Fig. S1**), but their ligand binding loops are engineered to specifically bind diverse molecular targets, including proteins, peptides and small molecules^28–30^. One of the targets, cytotoxic T-lymphocyte antigen 4 (CTLA-4), is involved in negative regulation of T-cell immune function and maintenance of immune homeostasis^31,32^, and represents an important target for cancer immunotherapy.

Here, we used AFM-SMFS to study the response of a non-antibody binding scaffold (lipocalin/anticalin) bound to its target (CTLA-4) and mechanically dissociated under a variety of pulling geometries. We systematically scanned the anchoring residue on anticalin from the N- to the C-terminus, targeting flexible loop regions located between secondary structural elements, and quantified the unbinding energy landscape for each anchor point. Our experimental approach was combined with molecular dynamics (MD) simulations, which helped us gain mechanistic insight into the unbinding pathways of the anticalin:CTLA-4 complex under different loading geometries. What emerges is a clear picture of geometrically optimal anchor points for protein receptor-ligands: pulling from central positions results in highly stable cooperative interactions that break at high forces, while pulling from the termini results in peeling-like behavior where anticalin loses contacts with CTLA-4 and breaks at low forces. These features amount to a molecular unbinding mechanism analogous to a suction cup being centrally pulled or peripherally peeled off a surface.

## Results

### Selection of anchor points, protein expression and AFM measurement setup

The anticalin targeting CTLA-4 was derived from human neutrophil gelatinase-associated lipocalin (NGAL). The structure of anticalin in complex with the extracellular domain of CTLA-4 is shown in **Fig. 1A** (PDB code 3BX7)^28^. CTLA-4 is a human T cell receptor with the C-terminus of the extracellular domain anchored to the cell surface. As shown in **Fig. S1**, the structure of anticalin is highly homologous to other previously reported lipocalin folds^33–37^, comprising a β-barrel formed by eight anti-parallel β-strands, flanked by helical regions at the N- and C-termini^38,39^. The two ends of the barrel are the open end, which is involved in ligand binding, and the closed end. Based on the protein structure, we selected eight anchor points on anticalin, scanning the sequence length through eight different pulling geometries. These anchor points were the N- and C-termini (residues 1 and 178), five residues located along flexible linkers connecting the β-strands (residues 21, 60, 87, 116 and 143), and one located within β-strand S2 (residues 55). All of the internal anchor points were located at the closed end of the barrel (**Fig. 1A and B**). N- and C-terminal anchor points were towards the open end of the barrel closer to the bound CTLA-4 ligand.

**Figure 1.**
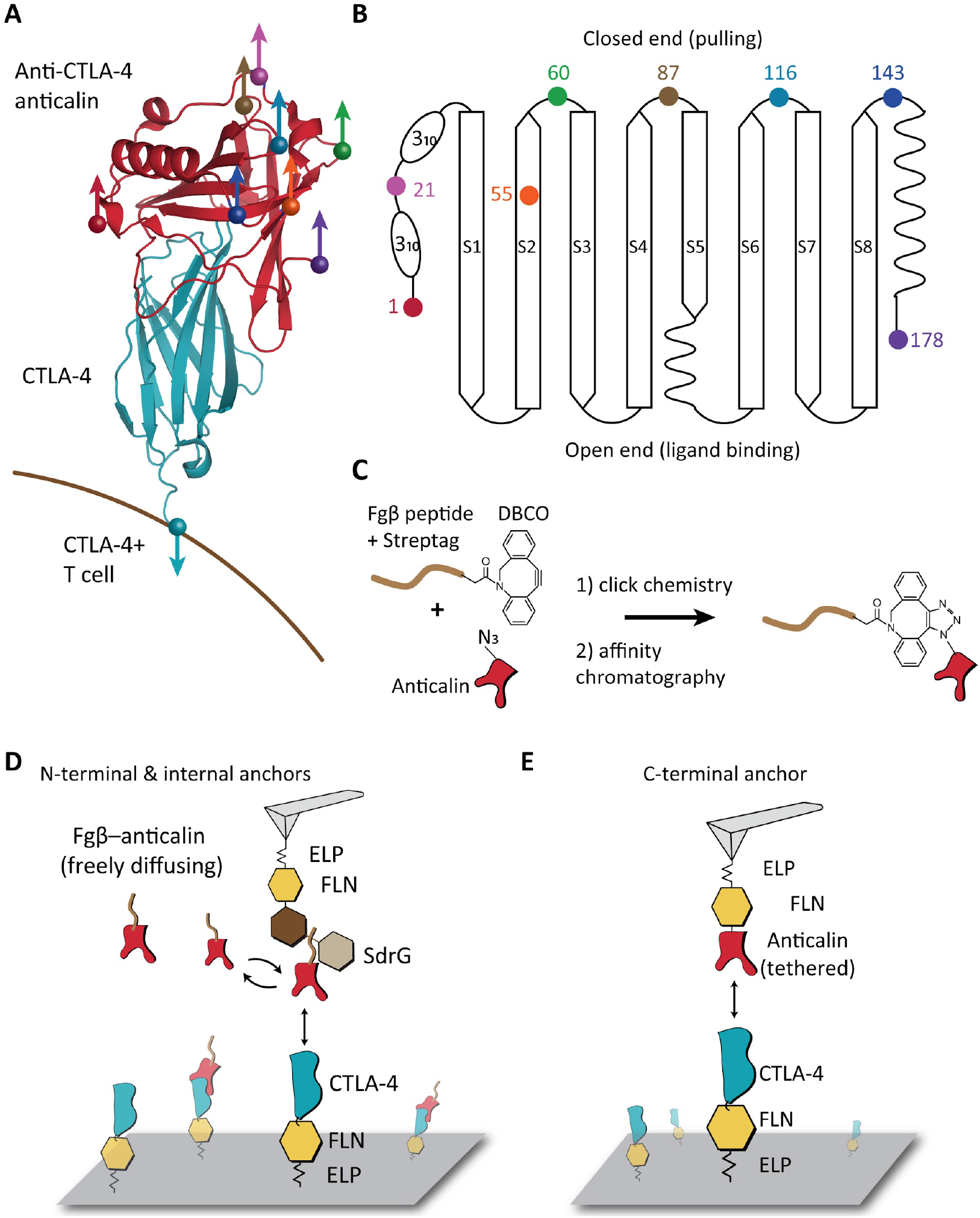
Anchor point selection and AFM-SMFS measurement setups. **A:** Structure of CTLA-4 in complex with anticalin (PDB code 3BX7). Anchor points on the anticalin are shown as colored spheres. The anchor point on CTLA-4 is fixed at the C-terminus, mimicking the natural tethering geometry on the cell surface. **B:** Anticalin has a central β-barrel, consisting of eight anti-parallel β-strands (S1-S8) connected by short flexible linkers. The anchor points shown as colored dots were chosen at the closed end of the β-barrel to avoid interference with the binding interface. **C:** Bioorthogonal conjugation of a fibrinogen β (Fgβ) peptide to the anticalin. The residue at the selected anchor point was replaced by p-azido-phenylalanine using amber suppression to introduce an azide group. The azide was covalently linked with a synthetic peptide comprising Fgβ-StrepTag and a C-terminal DBCO group using click chemistry. **D:** AFM measurement setup for testing N-terminal and internal anchor points with freely diffusing Fgβ-anticalin. Anticalin conjugated with Fgβ was added to the measurement buffer to final concentration of 1 µM. SdrG-FLN-ELP-ybbr was immobilized on the AFM tip and the ligand (CTLA-4-FLN-ELP-ybbr) was immobilized on the glass surface covalently via ybbr tag. **E:** AFM measurement setup with tethered anticalin for probing C-terminal anticalin anchor point. Anticalin-FLN-ELP-ybbr was immobilized on the cantilever and CTLA4-FLN-ELP-ybbr was immobilized on the glass surface.

Each anchor point on the anticalin molecule corresponds to a precisely defined pulling geometry in the AFM measurements. In order to attach the internal anchor points (residues 21, 55, 60, 87, 116 and 143) to the AFM tip, we combined non-canonical amino acid (NCAA) incorporation, click chemistry and a recently-reported AFM-SMFS setup with freely diffusing receptor molecules^11,23^. As shown in **Fig. 1C**, the residue at the selected anchor point was replaced by a p-azido-L-phenylalanine using amber suppression to introduce an azide group^40^ during protein translation in *E. coli*. A synthetic peptide comprising N-terminus fibrinogen β (Fgβ) peptide, Strep-tag and C-terminus dibenzocyclooctyne (DBCO) group was subsequently conjugated with the azide group on the anticalin using copper-free click chemistry^41^. The reaction product was purified with size-exclusion and Strep-trap columns to remove excess peptide and unreacted anticalin. Successful conjugation of the peptide increased the molecular weight of anticalin by 3 kD, as confirmed by SDS-PAGE and mass spectrometry (**Fig. S2**). Each anticalin with Fgβ clicked onto a given residue was expressed and purified, and measured in separate AFM experiments.

The AFM experimental setup with freely diffusing anticalin is shown in **Fig. 1D**. CTLA-4 was cloned to the N-terminus of a polyprotein, followed by an FLN unfolding fingerprint domain^42^, an elastin-like peptide (ELP) elastic linker, and a ybbr tag at the C-terminus. The ybbr tag facilitated site-specific and covalent surface immobilization of the polyprotein^43^ to the glass surface. The C-terminus of CTLA-4 was tethered to the glass surface, mimicking the natural pulling geometry on the cell surface. SD-repeat protein G (SdrG) from *S. epidermidis* which binds Fgβ was cloned into the polyprotein SdrG-FLN-ELP-ybbr and immobilized on the AFM tip.

The anticalin with Fgβ peptide conjugated to the selected anchor point was added to the measurement buffer to final concentration of ∼1 µM, which saturated immobilized CTLA-4 on the surface. SdrG on the cantilever can bind and unbind with Fgβ-anticalin in solution. A significant portion of the time, SdrG on the cantilever is free and when it approaches the surface, a 3-member complex forms, consisting of cantilever-borne SdrG bound to Fgβ-anticalin, which was itself in turn bound to CTLA-4. Alternatively, if SdrG was occupied by an Fgβ-anticalin molecule when the cantilever approached and indented the surface, mechanical contact can stimulate Fgβ ligand exchange, allowing the same 3-member complex to form. If no such exchange occurs, the AFM trace results in no specific interactions being recorded.

Despite the rapid on/off exchange of SdrG:Fgβ complexes at equilibrium, when placed under tension these complexes are extremely mechanically stable and can withstand forces as high as 2 nN ^44^. When retracting the cantilever the significantly weaker anticalin:CTLA-4 complex was therefore the first to break, leaving the Fgβ conjugated anticalin on the cantilever. Since the affinity between SdrG and Fgβ is mediocre (Kd ∼400 nM)^45^, the anticalin on the cantilever quickly exchanged with the freely diffusing anticalin molecules in the solution, preventing the AFM tip from clogging^11^. Tens of thousands of approach-retract cycles could be performed in this format repeatedly using a range of constant pulling speeds from 100 nm s^-1^ to 800 nm s^-1^ to build up large statistics. In this measurement format, it is important to note that the cantilever and surface molecules are always freshly probed, so the refoldability of the cantilever-borne molecules does not play a role.

For the N-terminal anchor point, anticalin was cloned and expressed with the Fgβ peptide at its N-terminus and measured using the freely diffusing setup. However, since the SdrG:Fgβ complex is itself directionally dependent, it was not possible to use Fgβ as a C-terminal anchor point. To anchor the anticalin from the C-terminus, we used an AFM setup with tethered anticalin, where a polyprotein containing anticalin at the N-terminus (anticalin-FLN-ELP-ybbr) was directly immobilized on the cantilever, as shown in **Fig. 1E**. The protocol for AFM cantilever approaching, dwelling and retracting was kept the same as for the setup with freely diffusing anticalin.

### Different pulling geometries gave rise to diverse unbinding energy profiles

In a typical AFM measurement (∼12 h), around 10,000 force-extension curves were recorded and transformed into contour length space using a freely rotating chain (FRC) elasticity model^46^. The curves were filtered based on the two-step unfolding pattern and 32 nm contour length increment of two FLN fingerprint domains^42^. In both AFM setups with freely diffusing or tethered anticalin, one FLN was on the cantilever and another was on the cover glass. We only analyzed the curves containing two FLN unfolding events to rule out nonspecific interactions and multiple parallel interactions between the tip and glass surface. Example force-extension curves of different pulling geometries are shown in **Figs. 2A and S3**. Intermediate unfolding events were observed in ∼9% of selected force curves, including all eight pulling geometries (**Fig. S3**). We aligned the contour length histograms of all the selected curves using cross-correlation analysis. The resulting superposition histogram (**Fig. S4**)^47,48^ showed contour length increments corresponding to the two FLN domains, which together added 64 nm of additional contour length to the system.

**Figure 2.**
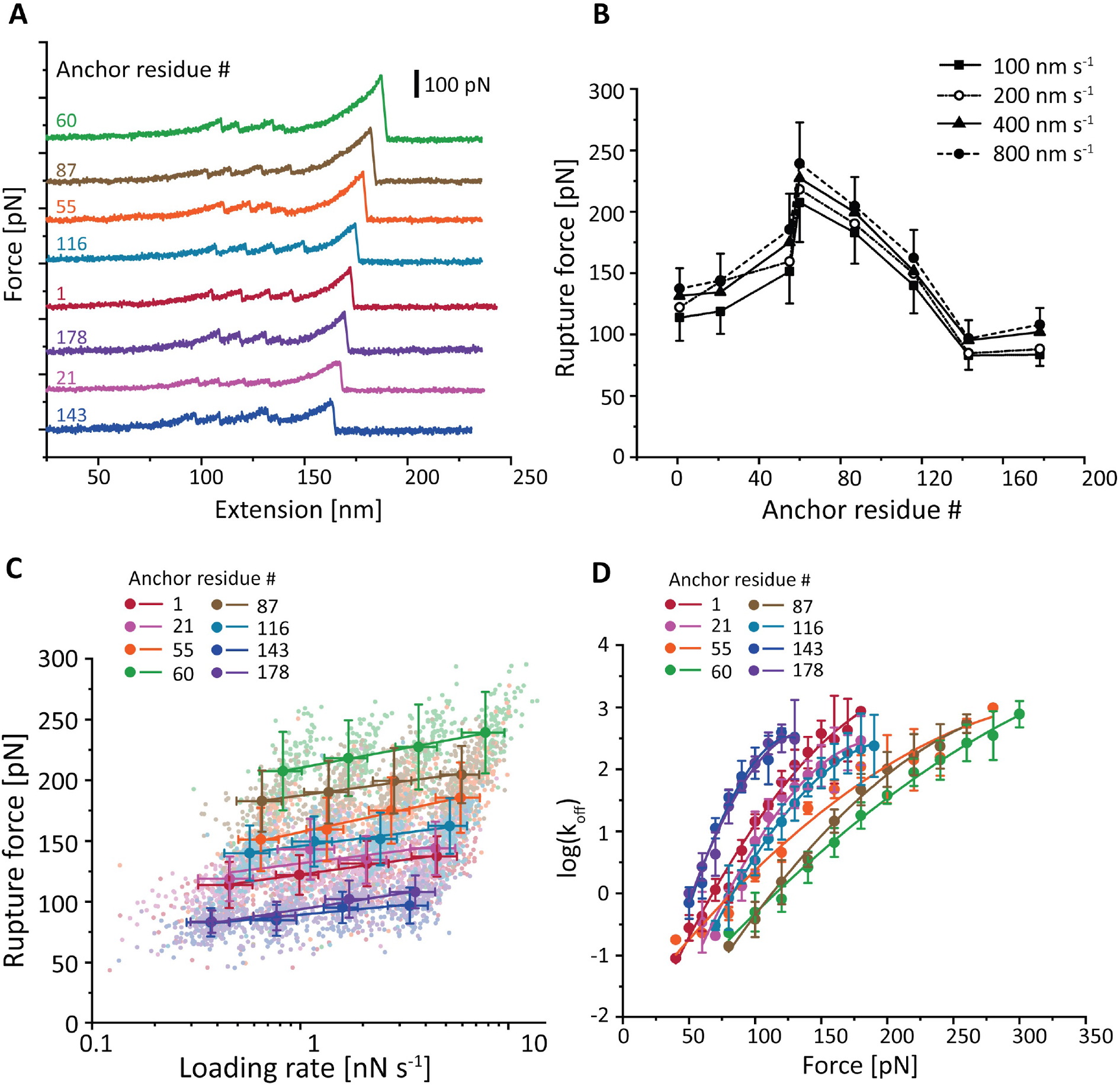
Dependency of anticalin:CTLA-4 complex stability on anticalin anchor position. **A:** Example AFM force-extension traces measured with eight different pulling geometries. Each trace shows unfolding of 2 FLN fingerprint domains and rupture of the anticalin:CTLA-4 complex for a given anchor residue on anticalin. **B:** Most probable rupture forces of the anticalin:CTLA-4 complex at various pulling speeds were plotted against the anchor residue number on anticalin. Error bars represent standard deviation of rupture forces measured at 100 nm s^-1^ (minus) and 800 nm s^-1^ (plus) pulling speeds. **C:** Most probable rupture forces measured at different pulling speeds were plotted against the logarithm of average loading rate and fitted linearly to extract the zero-force off rate *k*_*0*_ and distance to the transition state *Δx*^*‡*^. Error bars represent the standard deviation of rupture forces and loading rates. **D:** Force-dependent off rate of anticalin:CTLA-4 complex was plotted against force and fitted using Eq. 6 to extract *k*_*0*_, *Δx*^*‡*^ and *ΔG*^*‡*^. Error bars represent the standard deviation of off rates measured at four different pulling speeds.

Rupture forces of the anticalin:CTLA-4 complex for each Fgβ anchoring residue number were measured in separate AFM data collection runs at four different pulling speeds ranging from 100 to 800 nm s^-1^. The forces required to dissociate anticalin from CTLA-4 were plotted as histograms, as shown in **Fig. S5**. The histograms were fitted to extract the most probable rupture forces, which were plotted against the anchor residue number on anticalin (**Fig. 2B**). It is clear from the plot that the mechanical stability of the anticalin:CTLA-4 complex is highly dependent on the anticalin anchor residue number. N- and C-terminal pulling points were among the lowest in stability, rupturing at 100-125 pN. The stability significantly rose for anchor points located near the middle of the protein sequence, for example at residues 60 and 87. Peak stability (∼225 pN) was achieved when anticalin was anchored at residue 60, between β-strands S2 and S3.

We used Bell-Evans (BE)^49,50^ and Dudko-Hummer-Szabo (DHS)^51,52^ models to extract the unbinding energy profile parameters for each pulling geometry. As shown in **Fig. 2C**, the rupture forces measured at each pulling speed were plotted against the logarithm of loading rate, and the most probable rupture forces were linearly fitted against the average loading rate at a given pulling speed to extract the zero force off-rate (*k*_*0*_) and the distance to the energy barrier (*Δx*^*‡*^) using the BE model. We next used the DHS model (**Fig. 2D**) to transform the rupture force histograms into force-dependent off-rates (Eq. 4), and fitted the resulting plot using Eq. 6 to obtain the *k*_*0*_, *Δx*^*‡*^ and the height of the energy barrier (*ΔG*^*‡*^). The *k*_*0*_, *Δx*^*‡*^ and *ΔG*^*‡*^ values obtained using both models are listed in **Table 1**.

**Table 1.** Unbinding energy landscape parameters of the anticalin:CTLA-4 complex under different pulling geometries.

| Anchor point on anticalin | $k_0$ [$s^{-1}$] (DHS) | $k_0$ [$s^{-1}$] (BE) | $\Delta x^\ddagger$ [nm] (DHS) | $\Delta x^\ddagger$ [nm] (BE) | $\Delta G^\ddagger$ [$k_B T$] (DHS) |
| --- | --- | --- | --- | --- | --- |
| 1 | $(1.0 \pm 0.5) \times 10^{-3}$ | $(8 \pm 5) \times 10^{-4}$ | $0.50 \pm 0.05$ | $0.40 \pm 0.03$ | $16.8 \pm 0.6$ |
| 21 | $(7 \pm 9) \times 10^{-5}$ | $(1 \pm 7) \times 10^{-4}$ | $0.63 \pm 0.09$ | $0.4 \pm 0.2$ | $18 \pm 1$ |
| 55 | $(9 \pm 6) \times 10^{-3}$ | $(4 \pm 3) \times 10^{-3}$ | $0.27 \pm 0.04$ | $0.26 \pm 0.02$ | $13.9 \pm 0.9$ |
| 60 | $(2 \pm 1) \times 10^{-3}$ | $(2 \pm 1) \times 10^{-5}$ | $0.25 \pm 0.02$ | $0.30 \pm 0.01$ | $17 \pm 1$ |
| 87 | $(2 \pm 1) \times 10^{-4}$ | $(1 \pm 1) \times 10^{-6}$ | $0.38 \pm 0.03$ | $0.41 \pm 0.03$ | $17.8 \pm 0.4$ |
| 116 | $(1.0 \pm 0.3) \times 10^{-4}$ | $(2 \pm 4) \times 10^{-5}$ | $0.57 \pm 0.02$ | $0.44 \pm 0.07$ | $17.2 \pm 0.2$ |
| 143 | $(4 \pm 5) \times 10^{-5}$ | $(0.4 \pm 1) \times 10^{-3}$ | $1.0 \pm 0.1$ | $0.6 \pm 0.1$ | $18 \pm 1$ |
| 178 | $(2 \pm 2) \times 10^{-4}$ | $(3 \pm 2) \times 10^{-2}$ | $0.8 \pm 0.1$ | $0.36 \pm 0.04$ | $17 \pm 1$ |

It is clear from **Fig. 2** and **Table 1** that the complex crossed unbinding energy barriers with significantly different heights and shapes when pulled from different anchor points. Depending on the anchor residue number, the kinetic off-rate *k*_*off*_ at a given force can vary by 2-3 orders of magnitude (**Fig. 2D**, 100 pN). Both DHS and BE models extracted short *Δx*^*‡*^ values for anchor residue 60 (DHS: 0.25 ± 0.02 nm; BE 0.30 ± 0.01 nm) indicating a short steep energy barrier when tension was applied through the middle of the beta barrel on anticalin.

### Analysis with anisotropic network model

To help interpret our experimental observations, we applied an anisotropic network model (ANM) to calculate the effective spring constant between different pairs of residues^53,54^ in the protein complex. The ANM has been previously applied to several isolated protein domains including α-amylase inhibitor, green fluorescence protein (GFP), human ubiquitin and *E. coli* pyruvate dehydrogenase (PDH) E2lip3 domain, and the results were compared with MD simulations and experimental data^55,56^. Here we applied the ANM to the anticalin:CTLA-4 complex and derived effective spring constants between pairs of residues located on separate chains within the crystal structure. One residue was held fixed at the C-terminus of CTLA-4 while the other was scanned through all the residues on anticalin **(Fig. S6)**. To compare the predicted force constants from the ANM with the rupture forces and energy landscape parameters, we focused on the eight anchor residues on anticalin which were experimentally measured by AFM-SMFS **(Table S1)**. As shown in **Fig. S7**, the correlation between the experimentally measured rupture forces and the effective spring constant is not significant (*p* = 0.42). However, the *k*_*0*_ parameter derived using the DHS model, along with the *Δx*^*‡*^ value derived from the BE model were both statistically correlated (*p* < 0.05) with the effective spring constants between pairs of residues predicted by the ANM. The *ΔG*^*‡*^ parameter derived from the DHS model was weakly correlated (*p* = 0.07). Both *ΔG*^*‡*^ (DHS) and *Δx*^*‡*^ (BE and DHS) increased with increasing effective force constant, while *k*_*0*_ (DHS model) decreased with increasing effective force constant. The *k*_*0*_ calculated using the BE model, however, had negligible correlation with the effective force constant. We note that the unbinding energy profile measured for anticalin anchored from residue 55 was considered an outlier and excluded from the aforementioned correlation analysis. Residue 55 is the only anchor point located in a rigid region (β-strand) of anticalin. All other anchor residues are located at termini or within flexible loops. Due to its comparatively high structural rigidity, position 55 gave rise to the highest effective spring constant predicted by ANM, and resulted in a unique unbinding energy profile with a medium to low energy barrier and short *Δx*^*‡*^. Excluding residue 55, several of the experimentally determined energy landscape parameters were statistically correlated with the effective spring constant determined by the ANM approach. These findings provide evidence that ANM can be used to predict the anisotropic mechanical properties of protein-protein complex binding interfaces **(Figs. S6 and S7)**.

### Utilizing experimental data to improve hybrid all-atom and Gō-MARTINI descriptions of protein binding interfaces

Next we analyzed the anticalin:CTLA-4 system using a structure-based coarse grained model that combined the MARTINI force field^57,58^ with a Gō-like description of all native contacts that maintain secondary and tertiary structures in the protein. The latter model has been used in previous studies on mechanical stability of single protein domains^59–61^ and protein aggregates^62–64^. Combined Gō-MARTINI now allows analysis of large conformational changes in proteins, and the characterization of protein mechanical properties^65,66^. Gō-MARTINI simulations of anticalin:CTLA-4 dissociation under different pulling geometries at first failed to reproduce the rupture force profile as a function of residue position as obtained by SMFS (**Fig. S8**). This indicated an incomplete representation of the energetics at the interface, a known issue in MARTINI force fields^67^. Hence, an additional set of native contacts (NCs) obtained from the crystal structure (PDB code 3BX7) were included in the Gō-MARTINI model to better describe the binding interface between anticalin and CTLA-4 (**Fig. S8A**) using Gō contact map determination (http://pomalab.ippt.pan.pl/GoContactMap/). However, this minimal description of contacts was again insufficient to capture the experimental SMFS profiles (**Fig. S8B**). To provide qualitative agreement with experiments and remain consistent with a minimal representation of the binding interface dictated by structure, an additional native contact was introduced. This contact was chosen through a systematic process considering contacts established within a cutoff distance equal to 0.9 nm between any pair of Cα-atoms laying at the interface. Then, we performed pulling simulations for each of them to identify the one which improved the rupture profile. Thus, an additional contact between PRO102 (CTLA-4) and VAL66 (anticalin) was included in the contact map (CM). The updated CM improved the energetic description of the protein-protein interface and could be tuned to achieve simulations that reproduced the experimental rupture force profiles (**Fig. S8B**). In order to map the energetics of the all-atom MD simulation for the interface, the energy of the additional contact was tuned to reproduce atomistic energy in equilibrium. This step corrects the energy scale of the protein-protein interface via an energy parameter (ε_core_). Note that the standard native contact energy (ε_Gō-MARTINI_) is equal to 9.414 kJ mol^-1^ in the Gō-MARTINI model. The ε_core_ parameter was incrementally increased, and the system was evaluated via pulling simulations at different residue positions until the rupture force profile reached a qualitative agreement with the experiments (SMFS data from **Fig. 2B**). The corresponding value of ε_core_ that allowed this reconciliation was achieved at ε_core_ = 100 kJ mol^-1^ (see **Fig. S8B**). This result was validated in an ensemble (n=100) of different pulling simulations (see **Fig. S9**) for each anchor point. Although the optimal ε_core_ value is an order of magnitude larger than ε_Gō-MARTINI_, its value is bounded by the typical energy scales we found in all-atom MD simulation. The energetic decomposition for the anticalin:CTLA-4 complex allows us to verify the improvement given by the Gō-MARTINI description. **Table S2** shows the energy values for each protein chain and the protein-protein interface, which was defined by amino acid residues that form contacts between CTLA-4 and anticalin in the native structure. On average the energy at the interface in all-atom MD corresponded to 4% of total energy. The same decomposition for the Gō-MARTINI model assigned an average value of about 6% of the total energy. Note that our model retains the energy ratio for CTLA-4 and anticalin in good agreement with all-atom MD. The stability of the anticalin:CTLA-4 complex is mediated by the stability of the interface. The number of NCs was 189, 275 and 33 NCs for CTLA-4, anticalin and the protein-protein interface during 100 ns MD simulation.

### Calibrated Gō-MARTINI model provides molecular insights into deformation pathways

With the calibrated model established, we next analyzed the trajectories to help explain the observed differences in binding strength as a function of anchor point. The first analysis we carried out was to monitor the position of the center-of-mass (COM) of anticalin during the pulling simulations (see **Fig. 3A**). **Fig. 3BC** compares the COM position of anticalin at zero force and at the maximal force observed in the simulation (F_max_) for anticalin anchor residues 1 (low stability) and 60 (high stability). For anchor residue 1, at F_max_ the COM of anticalin was clearly shifted in the negative y-direction by ∼0.75 nm and in the negative x-direction by ∼0.2 nm. However, when pulling anticalin from residue 60, the anticalin COM stayed close to its original position, translating slightly in the positive y-direction (∼0.1 nm) and negative x-direction (∼0.2 nm). These differences suggest a scenario where pulling from the N-terminus results in a peeling-like behavior of anticalin off of CTLA-4, while pulling from position 60 results in a well-aligned system that cooperatively breaks without translating. Analysis of the xy translation of anticalin COM was carried out for each anchor residue under pulling simulations (**Fig. S10**). These plots show that COM translation behavior is distinct for each anchor point. The lowest stability anchor point tested experimentally (residue #143) also shows a broad distribution of translation values for anticalin COMs at F_max_ suggesting significant deformation of the complex and rearrangement under tension.

**Figure 3.**
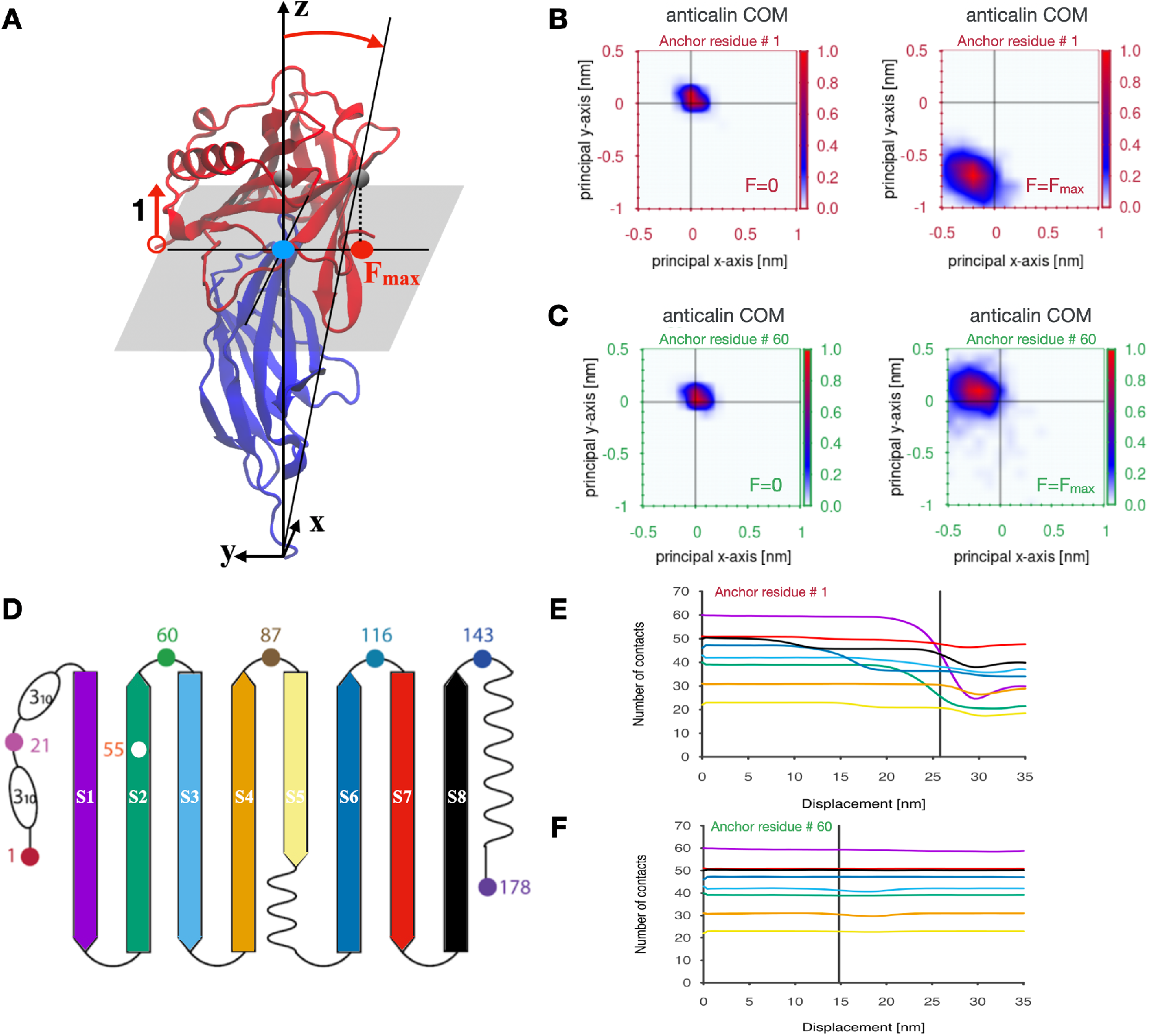
Molecular characterization of Gō-MARTINI trajectory for anticalin:CTLA-4 complex at different pulling geometries. **A:** The translation of anticalin COM for anchor residue 1 under a force applied along the z direction. The CTLA-4 was used as a reference system to define the normal plane. Blue circle denotes the starting anticalin COM position and the red one its translation along -y direction at F_max_. **B and C**: The relative translation of the anticalin COM with respect to CTLA-4 molecule for two anticalin anchor residues 1 **(B)** and 60 **(C)** at F = 0 and F = F_max_. Color bars indicate the probability of finding the COM in a given position along the X-Y plane which is perpendicular to the z direction of symmetry of the complex. **D:** The beta-sheet structure of the anticalin and its color representation. **E and F:** The intrachain native contact (NC) evolution for anticalin computed for each beta-sheet during the pulling process. Severe loss of contacts affects the anticalin for the pulling residue 1 **(E)**, whereas almost no loss of NC is reported for anchor residue 60 **(F)**. Color line is in agreement with the panel D.

The second MD analysis we carried out was to analyze the number of NCs (**Fig. 3EF and Fig. S11**) lost in different regions of anticalin when it was pulled from different directions. When pulling from residue 1, NCs were steadily lost in N-terminal beta strands S1, S2, S3 as well as S6 prior to rupture. However, when pulling from residue 60, few to no intramolecular NCs were lost in the anticalin (**Fig. 3F**). The anticalin COM shift is more pronounced when several anticalin beta-sheets lose some of the stabilizing NCs (**Fig. S11**). Our analysis of the intrachain NCs suggest that breaking NCs in the beta strands makes the anticalin more flexible and its COM samples new positions through different pathways. Furthermore, pathways involving partial unfolding processes and severe loss of NCs were observed for anchor residues 1 and 21 (**Fig. S11**). The NCs on the binding interface also behave differently depending on the pulling geometry (**Table S3 and Fig. S12**). The interface NCs were lost at different rates with different pulling geometries and the number of remaining interface NCs at F_max_ (immediately prior to rupture) varies across the simulations and shows positive correlation (p<0.05) with the rupture force measured both *in vitro* and *in silico* (**Fig. S13**). In addition, a few non-native contacts (about five, see **Fig. S12**) are established after the rupture. However, the new protein-protein interactions established during the dissociation pathway are not strong enough to maintain the mechanical stability of anticalin:CTLA-4 complex. Based on the simulation analyses, we conclude that the persistence of the original set of interface NCs, the translation of the anticalin COM and the loss of beta strand structure explain the geometric dependency of the mechanical properties of the complex.

## Discussion

Here, we reported a novel AFM-SMFS experimental method to covalently click a mechanostable peptide handle to internal residues of the non-antibody scaffold anticalin/lipocalin and to measure the rupture forces of anticalin:CTLA-4 complexes using an AFM setup with freely diffusing molecules. Using this method, we observed how the anticalin:CTLA-4 complex responds to external forces applied from different directions and found that the mechanical stability of the complex is highly dependent on the pulling geometry. When pulling from anticalin residue 60, the complex could withstand a high force up to 200 pN, which is ∼100% higher than the least mechanically stable pulling geometry (residue 143). To confirm that the different constructs have similar equilibrium affinity, we measured the dissociation constant between CTLA-4 and three anticlain mutants using microscale thermophoresis (MST). As shown in **Table S4**, although the anticalin E60AzF and E143AzF mutants have distinct responses to forces when pulled from different geometries, they have similar affinity towards CTLA-4 at equilibrium. The exception is the I55AzF mutant, which has a slightly lower affinity with CTLA-4. The anticalin mutants were conjugated with a bulky molecule using DBCO-azide reaction in both SMFS (with Fgβ peptide) and MST (with AF647 dye) measurements. Since residue 55 is on a β-strand, which is a part of the anticalin β-barrel structure, the mutation to pAzF and conjugation of peptide or fluorescent dye may have slightly destabilized the anticalin:CTLA-4 complex. Therefore, the I55AzF mutant gave rise to a lower energy barrier in SMFS measurement (see **Table 1** and **Fig. S7**) as well as lower equilibrium affinity in bulk experiments compared to other anticalin constructs.

**Fig. 4AB** illustrates the complex dissociation energy landscape. In the absence of force, the complex has the same free energy in different pulling geometry scenarios. When external force is applied to the complex from different anchor points, the complex unbinds through different pathways. These pathways correspond to unbinding energy barriers with different shapes and heights, giving rise to distinct responses to forces. It is worth noting that the energy barrier of the most mechanostable pulling geometry (anchor residue 60) is not the highest, but its exceptionally short *Δx*^*‡*^ contributed to the high resistance to external force. On the contrary, although the pulling geometry with anchor residue 143 has the highest energy barrier, the long *Δx*^*‡*^ made it the least mechanostable pulling geometry. Another interesting pulling geometry, anchor residue 55, has a unique energy landscape with the lowest energy barrier and a short *Δx*^*‡*^, giving rise to a mediocre rupture force among all pulling geometries. Therefore, the mechanical stability of the complex is determined by an interplay between the height and the shape of unbinding energy barriers.

**Figure 4.**
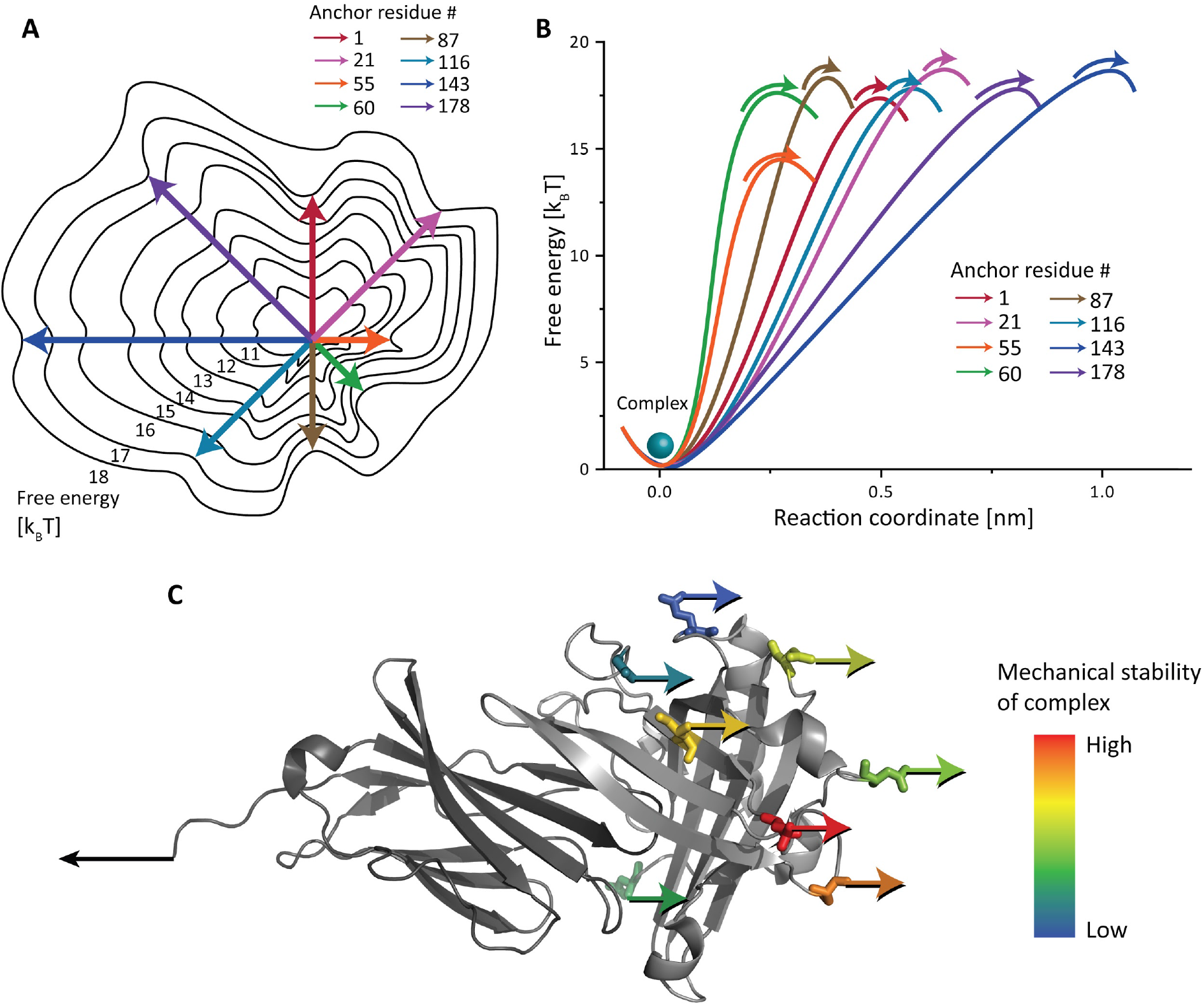
Depictions of anticalin:CTLA-4 complex unbinding energy landscape as a function of molecular pulling geometry. **A:** Energy landscape depiction where anchor point residues are represented as cardinal and ordinal directions of a compass. Under a constrained pulling geometry, the complex is forced to traverse different unbinding pathways across the energy landscape. These different paths give rise to energy barriers with diverse heights and shapes. **B:** 1D depiction of unbinding energy barrier heights and positions calculated using the DHS model for each pulling geometry (see **Table 1**). **C:** Anchor points on the anticalin colored based on mechanical stability of the complex pulled through that position. The most and least mechanostable anchor points on the anticalin are residues 60 (red) and 143 (dark blue), respectively.

We further used the anisotropic network model (ANM) to calculate the force constant between the eight anchor residues on the anticalin and the C-terminal anchor point of CTLA-4. While the effective force constant has very weak correlation with the rupture force, it is correlated with many of the energy profile parameters calculated using both Bell-Evans and Dudko-Hummer-Szabo models. However, residue 55 of anticalin, which is located on a rigid β strand while all the other anticalin anchor points are at the termini or in flexible loops, is an outlier in all of these correlations. This indicates that the AMN approach has certain limitations and additional factors, for example secondary structure, should be accounted for when applying this method.

The computational approach for investigating the nanomechanical stability of the anticalin:CTLA-4 complex under different pulling directions was parametrized by tuning the interface energy. This rendered the Gō-MARTINI approach a very predictable model for the study of large conformational changes of protein complexes at much cheaper computational cost than in regular SMD simulation allowing larger sampling of pathways. This model not only allowed the reproduction of the phenomenological one-to-one experimental SMFS profile for anticalin:CTLA-4 complex, but also unveiled microscopic pathways and mechanistic explanations of the higher mechanical stability observed when pulling anticalin at residue 60. Our computational study explained this stability in terms of translations of anticalin COM, and loss of NCs in anticalin. The simulations show that certain pulling directions can partially unfold the anticalin and destabilize the interface while other positions maintain the close contact of the proteins at the interface.

The anticalin against CTLA-4 shares very high structural similarity with other previously reported engineered anticalins (**Fig. S1**). In the future, it will be interesting to see if the mechanically optimal pulling point (residue 60) is generalizable to other anticalin:ligand complexes. Our research demonstrated that the mechanical stability of protein-ligand systems can be tuned by precisely controlling the loading geometry, without changing equilibrium binding properties. This suggests a new paradigm for affinity maturation of non-antibody scaffolds by correctly choosing the anchor points. Such an approach could be particularly beneficial for targeting therapeutic nanoparticles or imaging probes which can exert shear forces onto binding interfaces.

## Methods

### Expression and purification of Fgβ-anticalin and anticalin-ddFLN4-ELP-ybbr

The constructs were cloned in a pET28a vector, which was used to transform NiCo21(DE3) competent cells (New England Biolabs, Ipswich, MA, USA). The cells were grown in terrific broth (TB) medium containing 50 µg mL^-1^ kanamycin at 37 °C until OD reached ∼ 0.6. The protein expression was induced by adding isopropyl β-D-1-thiogalactopyranoside (IPTG) to a final concentration of 0.5 mM and incubating at 20 °C overnight. The cells were subsequently pelleted and lysed using sonication. The cell lysate was loaded to a His-trap column (GE Healthcare, IL, USA), washed with phosphate buffered saline (PBS, 137 mM NaCl, 2.7 mM KCl, 10 mM Na_2_HPO_4_ and 2 mM KH_2_PO_4_, pH = 7.4) with 20 mM imidazole and eluted with PBS buffer supplemented with 500 mM imidazole. The eluate was further purified with Superdex Increase 200 10/300 GL size-exclusion column (GE Healthcare).

### Expression and purification of CTLA-4-ddFLN4-ELP-ybbr

CTLA-4 contains two disulfide bonds and therefore we used a system to facilitate cytoplasmic disulfide bond formation in E.coli (CyDisCo) to express the CTLA-4-ddFLN4-ELP-ybbr fusion protein. We co-transformed NiCo21(DE3) competent cells with pET28a vector carrying CTLA-4-ddFLN4-ELP-ybbr and pMJS205 vector carrying sulfhydryl oxidase Erv1p and disulfide bond isomerase PDI^68^. The transformed cells were grown in TB medium containing 50 µg mL^-1^ kanamycin and 34 µg mL^-1^ chloramphenicol at 37 °C until OD reached ∼ 0.8. The expression of Erv1p, PDI and CTLA-4-ddFLN4-ELP-ybbr was induced by adding 0.5 mM IPTG, followed by incubating at 20 °C overnight. The CTLA-4-ddFLN4-ELP-ybbr protein was purified using the same procedure as the Fgβ-anticalin construct.

### Amber suppression

Anticalin constructs with internal anchor points were expressed using an amber suppression system. The wild-type anticalin was cloned to a pET28a vector and the codon encoding the amino acid at the selected anchor point was mutated to an amber codon (TAG). NiCo21(DE3) competent cells were co-transformed using the expression vector and plasmid pEVOL-pAzF (addgene #31186). Cells were grown in 500 mL LB medium supplemented with 50 µg mL^-1^ kanamycin and 25 µg mL^-1^ chloramphenicol at 37 °C until OD reached 0.8, washed with 500 mL M9 minimal medium and transferred to 250 mL M9 medium supplemented with 50 µg mL^-1^ kanamycin, 25 µg mL^-1^ chloramphenicol, 0.2 mg ml^-1^ p-azido-l-phenylalanine (pAzF) and 0.02% arabinose. The culture was incubated at 37 °C for 1 h and was supplemented with 1 mM IPTG, followed by incubation at 20 °C overnight. The cells were lysed and the protein was purified using the same procedure as the wild-type Fgβ-anticalin construct.

### Conjugation of Fgβ peptide and anticalin

5x molar excess of Fgβ-StrepTag-DBCO peptide (JPT Peptide Technologies GmbH, Berlin, Germany) was added to AzF-incorporated anticalin. The mixture was incubated at room temperature with shaking for 1 h, followed by incubation at 4 °C overnight. The reaction mixture was purified with size-exclusion column to remove the excess peptide. The protein purified with SEC was loaded to a Strep-Trap column (GE Healthcare) and eluted using PBS buffer with 2.5 mM desthiobiotin to remove unreacted anticalin.

### Microscale thermophoresis measurements

The anticalin I55AzF, E60AzF and E143AzF mutants were labeled with AF647 fluorescent dye (Jena Bioscience, Jena, Germany) by mixing the protein with 5x molar excess of DBCO-AF647, incubating at room temperature for 1 h, and incubating at 4 °C overnight. The labeled proteins were separated from unreacted free dye molecules using size-exclusion column.

The titration samples were prepared by mixing 10 nM labeled anticalin with a series of unlabeled CTLA-4 with different concentrations ranging from 110 pM to 3.6 µM. The microscale thermophoresis (MST) traces of each titration sample were measured using a Nanotemper Monolith NT.115 (NanoTemper Technologies GmbH, Munich, Germany). The temperature-related fluorescence intensity change was recorded for each sample and plotted against the concentration of CTLA-4 to derive the dissociation constant between anticalin and CTLA-4.

### Surface chemistry for AFM measurements

Cover glasses and Biolever mini (Olympus) AFM cantilevers were amino-silanized with (3-aminopropyl)-dimethyl-ethoxysilane (APDMES, ABCR GmbH, Karlsruhe Germany). The silanized surfaces were incubated in 10 mg/mL sulfosuccinimidyl 4-(N-maleimidomethyl)cyclohexane-1-carboxylate (sulfo-SMCC, Thermo Fisher Scientific, Waltham, MA, USA) at room temperature for 30 min, followed by incubation in 20 mM coenzyme A (CoA, Sigma-Aldrich, St. Louis, MO, USA) at room temperature for 2 h. The cover glasses and cantilevers were extensively washed with ddH_2_O after each incubation step. The CoA-coated cover classes and cantilevers were incubated in CoA-ybbr reaction mixture, consisting of ∼40 µM ybbr-tagged protein, 5 µM Sfp (phosphopantetheinyl transferase) enzyme and 20 mM MgCl_2_, at room temperature for 2 h and subsequently washed with PBS buffer and stored in PBS until further use.

### AFM measurements

AFM-SMFS measurements were performed on a Force Robot AFM (Bruker, Billerica, MA, USA). The contact-free method was used to calibrate the cantilever spring constants (ranging from 0.02 N m^-1^ to 0.14 N m^-1^) and detector sensitivity. The cantilever and cover glass were submerged in PBS buffer. In the measurements using freely diffusing anticalin, ∼ 1 µM Fgβ-conjugated anticalin was added to the measurement buffer. The cantilever was approached to the glass surface, dwelled for 200 ms and retracted at a constant speed ranging from 100 nm s^-1^ to 800 nm s^-1^ (100 nm s^-1^, 200 nm s^-1^, 400 nm s^-1^ and 800 nm s^-1^). The x-y position of the stage was moved by 100 nm after each approach-retraction cycle to probe a new molecule on the surface.

### AFM data analysis

The recorded force-extension curves were transformed to contour length using the freely rotating chain (FRC) model. This model assumes that the polymer chain consists of bonds with length *b* and fixed angle *ϒ*, as described by Eq. 1^46^.

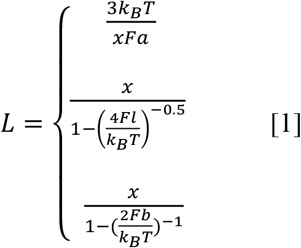

where *k*_*B*_ is the Boltzmann constant, *T* is temperature, 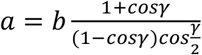 is the Kuhn length and 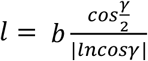 is the persistence length.

The force-extension curves were subsequently filtered based on the 64 nm contour length increment from the unfolding two FLN fingerprint domains. The final rupture forces and force loading rates were extracted from the selected force-extension curves. The rupture forces measured at different pulling speeds were plotted in histograms and the rupture force distribution *p*(*F*) was fitted with Bell-Evans model (Eq. 2)^50^ to extract the most probable rupture force.

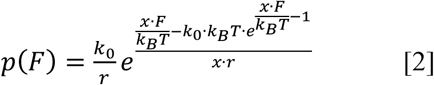

where *k*_*0*_ is the off rate at zero force, *r* is the loading rate, *Δx*^*‡*^ is the distance to the energy barrier, *k*_*B*_ is the Boltzmann constant and *T* is temperature.

The rupture force was plotted against the logarithm of loading rate and the most probable rupture force at each pulling speed was fitted against the average loading rate using a linear model to extract *k*_*0*_ and *Δx*^*‡*^, as described by the Bell-Evans model (Eq. 3)^49,50^:

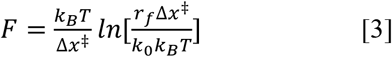

where k_B_ is the Boltzmann constant and T is the temperature.

In addition to the Bell-Evans model, we used the Dudko-Hummer-Szabo model to calculate the energy barrier parameters^51,52^. All rupture force histograms were plotted with equal binwidth *ΔF* = 20 pN. Each bin of the rupture force histogram was transformed into force-dependent off rate using Eq. 4:

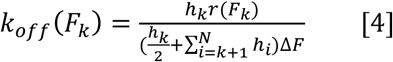

where *k*_*off*_(*F*_*k*_) is the force-dependent off rate at the average rupture force of the k_th_ bin, *r*(*F*_*k*_) is the average loading rate of the k_th_ bin, and *h*_*k*_ is the height of the k_th_ bin, which is calculated using Eq. 5:

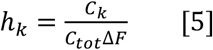

where *C*_*k*_ is the number of counts in the k_th_ bin and *C*_*tot*_ is the total number of counts in the histogram.

The calculated force-dependent *k*_*off*_(*F*) was plotted against force and fitted using Eq. 6:

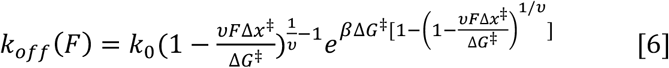

where *k*_*0*_ is the zero-force off rate, *Δx*^*‡*^ is the distance to the energy barrier, *ΔG*^*‡*^ is the height of the energy barrier in the absence of force, *β*^-1^*=k*_*B*_*T*, and *υ* = 0.5, which assumes a cusp shape of the free energy surface.

### Calculation of effective spring constant using anisotropic network model

The effective spring constants were calculated using the MechStiff function of the ProDy Python package^56,69^. The Python code (see Supplementary Note 2) was adapted from the mechanical stiffness calculations tutorial of ProDy, available at http://prody.csb.pitt.edu/tutorials/mech_stiff/sm.html. The first five residues of anticalin are missing in the protein structure. Therefore the residue 6 of anticalin was considered as the N-terminus when calculating the force constant between the C-terminus of CTLA-4 and the anticalin residues.

### Computational Modelling

The crystallographic structure of the anticalin:CTLA-4 complex was taken from the deposited structure (PDB code 3BX7). Several missing residues were reconstructed using MODELLER^70^ and UCSF Chimera^71^. In order to obtain a well-defined native contact map for the Gō-MARTINI simulation we employed an all-atom MD description in explicit water of the anticalin:CTLA-4 complex. The system was simulated by the CHARMM36c force field in GROMACS software^72^. The long-range electrostatic interactions were evaluated using the particle mesh Ewald (PME) method^73^ with a grid size of less than 0.12 nm. A time step equal to 2 fs was used for integration of the potential function. The system was solvated with TIP3P water molecules and Na+ and Cl- ions were added to achieve the concentration of 150 mM and neutralize the whole system. Pre-equilibration steps were employed by first running an energy minimization step followed by equilibration in NVT ensemble at 300 K and final equilibration step was done in NPT ensemble. The temperature and the pressure of the system were set to 300 K and 1 bar using V-Rescale thermostat^74^ and Berendsen barostat coupling methods. The production run was carried out at the same thermodynamics parameters of temperature and pressure using V-Rescale and Parrinello-Rahman algorithms respectively for 100 ns. The molecular trajectory was used to calculate the native contact map. In total we got 1000 maps based on the rCSU+OV contact map determination protocol. Only contacts with high frequencies (i.e. larger than 0.9) were considered. The set of contacts chosen avoid placing contact in flexible loops or other highly flexible regions in each protein chain.

The atomistic system was mapped onto a coarse-grained resolution via the MARTINI approach^57^ and the secondary and tertiary structure were retained by the Gō-MARTINI interactions^65^ with an effective depth (ε) of Lennard-Jones (LJ) potential equal to 9.414 kJ mol^-1^ and the corresponding σ_*ij*_ of the LJ potential is determined by the following relationship, σ_*ij*_ = *d*/2^1/6^, with *d* being the Cα–Cα distance between a pair of residues that form a native contact. The set of native contacts or Gō-MARTINI interactions were determined through our local server (http://pomalab.ippt.pan.pl/GoContactMap/). From the all-atom simulation in equilibrium we obtained the following set of contacts for each protein region 189, 275 and 33 for CTLA-4, anticalin, and anticalin:CTLA-4 interface, respectively. The coarse-grained model was pre-equilibrated similarly to the all-atom MD model.

### Characterization of the mechanical stability and unbinding process of proteins by Gō-MARTINI

The nanomechanics of the protein complexes were investigated through a Gō-MARTINI approach that is based on the combination of the MARTINI model^57^ and the Gō-like model^59,61,64^. As a result the Gō-MARTINI model has been used to sample large conformational changes in proteins and their aggregates^65^. The former method offers high computational efficiency, while the latter gives an accurate (as much as all-atom MD) energetic description of the process as long as an accurate native contact map is determined. In order to quantify the differences in mechanical stability for the anticalin:CTLA-4 complex, we performed stretching simulations. The pulling direction was chosen along the end-to-end vector connecting the Cα-atom of the CTLA-4 C-terminus and an anchor point in anticalin that was pulled along this vector. Moreover, additional beads have to be attached to those Cα-atoms with the spring constant being 37.6 kJ mol^-1^ nm^-2^, which is a reasonable value of the AFM cantilever stiffness in protein stretching studies. The typical time scale in the pulling simulations was on the order of 60 ns with a velocity of 5×10^7^ nm s^-1^, with rupture of the complex generally occurring at an earlier stage. Although this value is still far from the experimental values of cantilever velocities (100-800 nm s^-1^), it represents a significant computational improvement compared to all-atom SMD whose typical pulling speed is about an order of magnitude larger. In practice, the Gō-MARTINI model allows characterization of the process occurring in the SMFS experiment at almost the same resolution as conventional all-atom MD.

In AFM-SMFS, multiple proteins or protein complexes can exhibit several peaks in the force-distance profile, which signal the partial unfolding of one protein domain or component. Because of the limitations on time resolution and cantilever sensitivity, intermediate unfolding states are often missed in the experimental datasets. However, in the case of our computational model we can observe these intermediate states with a better resolution and assign to each of them a force peak. The largest of these force peaks, F_max_, defines the characteristic largest rupture force in the system before dissociation or rupture of the protein-protein interface. Also, we can carry out an analysis of the presence of native contact at the rupture force. A contact will be present if the native distance, *d*, between two Cα atoms is less than 1.5 σ_*ij*_.The analysis of the mechanical stability required a set of 100 independent trajectories. To fine-tune our data and reproduce the experimental AFM-SMFS rupture profile (see **Fig. 2B**), the strength of the contact between residues VAL66 and PRO102 in anticalin and CTLA-4 was increased. The energy scale (ε_core_) that optimized the agreement with the SMFS experiment was in the range of 100 kJ mol^-1^ and was bounded by protein-protein interface energy. The determination of non-native contacts in Gō-MARTINI simulations was constructed based on the overlaps of the enlarged VdW radii centered on Cα atoms. VdW parameters were taken according to our previous work^59^. If the distance of a pair of Cα atoms is smaller than the sum of their enlarged VdW radius, then we count the pair as a non-native contact. In order to characterize the unbinding process in Gō-MARTINI simulations we devised a reference system using the CTLA-4 as no large fluctuations were observed for that molecule. Then, anticalin was aligned with respect to CTLA-4. This allowed us to calculate the main axis of symmetry along the pulling z-direction and define the plane perpendicular to it in the x-y coordinates. For each snapshot in the molecular trajectory the center-of-mass of the anticalin was monitored and its changes were projected onto this plane (see **Fig. S10**).

## Acknowledgements

This work was supported by the University of Basel, ETH Zurich, an ERC Starting Grant (MMA-715207), the NCCR in Molecular Systems Engineering, and the Swiss National Science Foundation (Project 200021_175478). The authors thank Peter Schultz for providing the pEVOL-pAzF plasmid and Lloyd Ruddock for providing the pMJS205 plasmid. The authors thank Timothy Sharpe and the Biophysics Facility of Biozentrum, University of Basel for the help with affinity measurements using MST. A.B.P. acknowledges financial support from the National Science Centre, Poland, under grant No. 2017/26/D/NZ1/00466, also gratefully acknowledge the computing provided by the Jülich Supercomputing Centre on the supercomputer JURECA at Forschungszentrum Jülich and additional computer resources were supported by the PL-GRID infrastructure.

## Supporting Information Available

Figures S1-S13, Tables S1-S4 and Supplementary Notes 1 and 2

