## Supplementary Information for "Optimizing mechanostable anchor points of engineered lipocalin in complex with CTLA-4"

### Table of contents

**Supplementary Figures**

Figure S1. Structure and sequence alignments between anticalin targeting CTLA-4 and other lipocalin folds

Figure S2. Successful conjugation of Fgβ to anticalin demonstrated by SDS-PAGE and MS

Figure S3. Example force-extension curves with intermediate unfolding steps

Figure S4. Combined contour length histograms measured at different pulling geometries

Figure S5. Rupture force histograms of the anticalin:CTLA-4 complex under different pulling geometries

Figure S6. Effective force constant between the C-terminus of CTLA-4 and all anticalin residues

Figure S7. Correlations between rupture force or unbinding energy profile and effective spring constant between anchor points

Figure S8. Molecular dynamics (MD) simulations and in silico force spectroscopy

Figure S9. Example force-extension curves and rupture force histograms of anticalin:CTLA-4 complex at different pulling geometries in Gō-MARTINI simulation

Figure S10. Relative motion of the anticalin COM during Gō-MARTINI stretching simulations of the anticalin:CTLA-4 complex

Figure S11. Evolution of the anticalin intrachain native contacts (NC) during Gō-MARTINI stretching simulations of anticalin:CTLA-4 complex

Figure S12. Profile of native (open circles) and non-native (solid line) interface contacts during Gō-MARTINI stretching simulations of anticalin:CTLA-4 complex

Figure S13. Rupture forces vs. number of remaining NC at complex rupture

**Supplementary Tables**

Table S1. Effective spring constant between the C terminus of CTLA-4 and different anchor points on anticalin

Table S2. Non-bonded (VdW and coulomb interactions) energy contribution for each protein chain in the anticalin:CTLA-4 complex and protein-protein interface at different levels of representation

Table S3. Statistics of the total number of native contacts (NC) present in each protein component of anticalin:CTLA-4 complex at different pulling geometries.

Table S4. Dissociation constant between CTLA-4 and anticalin mutants

**Supplementary Notes**

Supplementary Note 1 Amino acid sequences and Addgene accession codes

Supplementary Note 2 Python code used to calculate the effective force constants

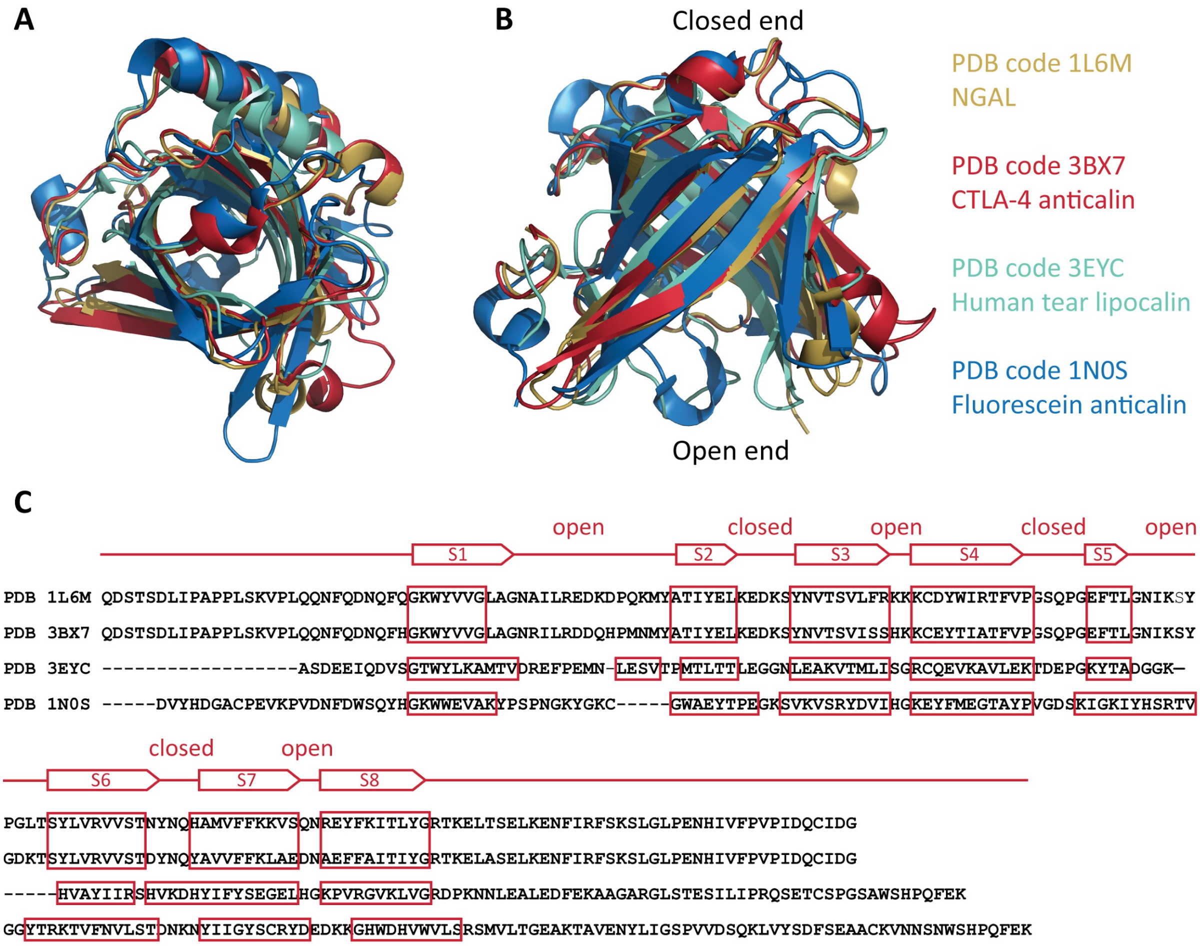


**Figure S1. Structure and sequence alignments between anticalin targeting CTLA-4 and other lipocalin folds. A and B:** The structure of anticalin targeting human CTLA-4 (PDB 3BX7) was aligned with neutrophil gelatinase-associated lipocalin (NGAL, PDB 1L6M), human tear lipocalin (PDB 3EYC) and an anticalin binding fluorescein (PDB 1N0S). The alignment is shown in top view (panel A, from the closed end) and side view (panel B). Lipocalins share the same β-barrel structure formed by eight β-strands. **C:** Sequence alignment of lipocalins. The anti-(CTLA-4) anticalin was derived from NGAL and has a very high sequence homology (84% identity) with NGAL. The different residues are mainly on the open end, which was engineered to bind CTLA-4. The human tear lipocalin and fluorescein anticalin, which was derived from bilin-binding protein (BBP), have very low sequence homology but high structure homology with CTLA-4 anticalin and NGAL.

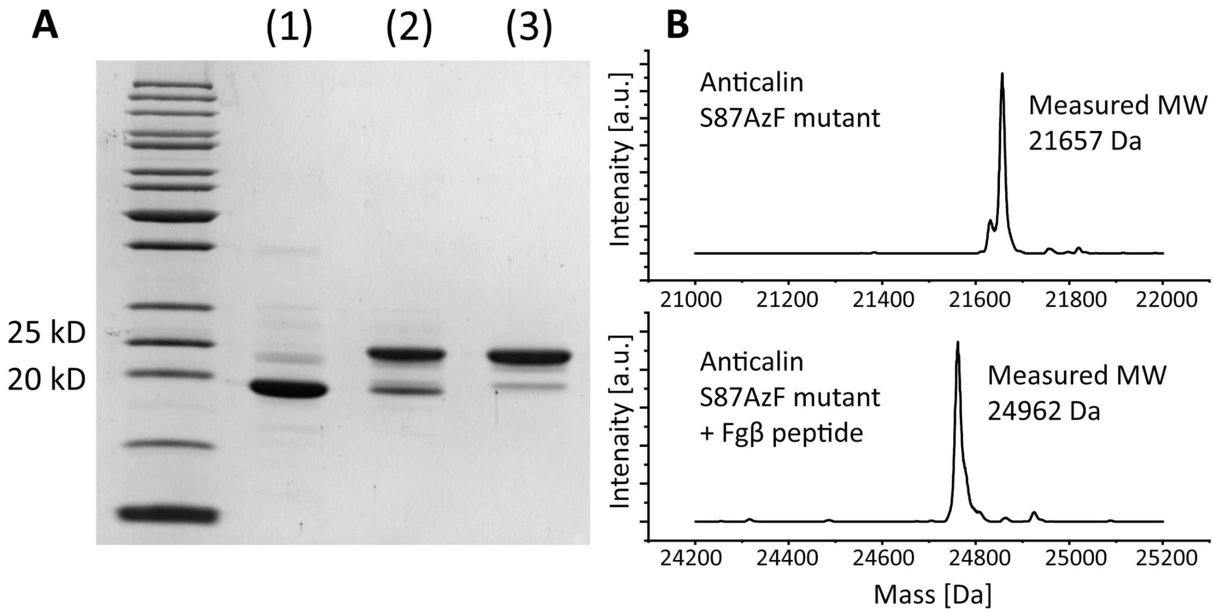


**Figure S2. Successful conjugation of Fgβ to anticalin demonstrated by SDS-PAGE and MS. A:** SDS-PAGE analysis of anticalin S87AzF mutant protein (lane 1) and the product of the conjugation reaction with Fgβ-StrepTag-DBCO peptide before (lane 2) and after (lane 3) Strep-Trap column purification. Successful conjugation of the peptide increased the protein molecular weight by ~3 kD. Unreacted S87AzF protein was mostly removed by the Strep-Trap column. **B:** Mass spectrometry measurements on anticalin S87AzF mutant before and after conjugation with Fgβ-StrepTag-DBCO. The theoretical molecular weight of S87AzF mutant is 21,659 Da. After conjugation with the Fgβ peptide, the molecular weight increased by 3,305 Da, exactly matching the molecular weight of the synthetic Fgβ-StrepTag-DBCO peptide.

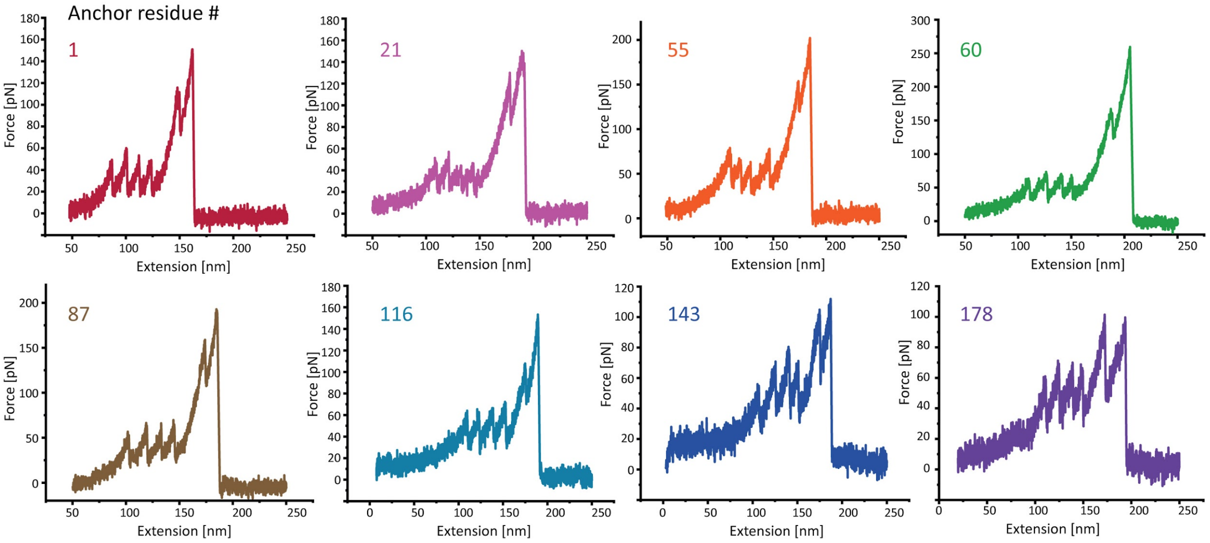


**Figure S3. Example force-extension curves with intermediate unfolding steps.** Around 9% of the observed curves have intermediate unfolding events prior to complex rupture. Intermediate unfolding events were observed in all the eight pulling geometries suggesting the unfolding is attributable to partial unfolding of CTLA-4.

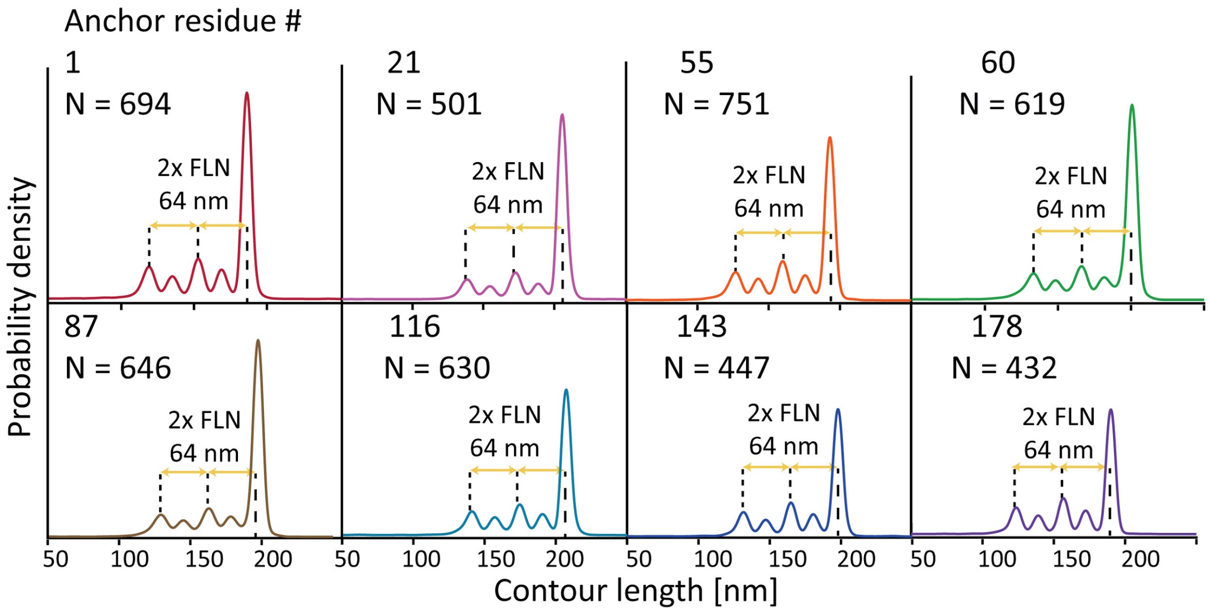


**Figure S4. Combined contour length histograms measured at different pulling geometries.** All histograms show the unfolding of two FLN fingerprint domains, giving rise to 64 nm in total contour length prior to dissociation of the anticalin:CTLA-4 complex.

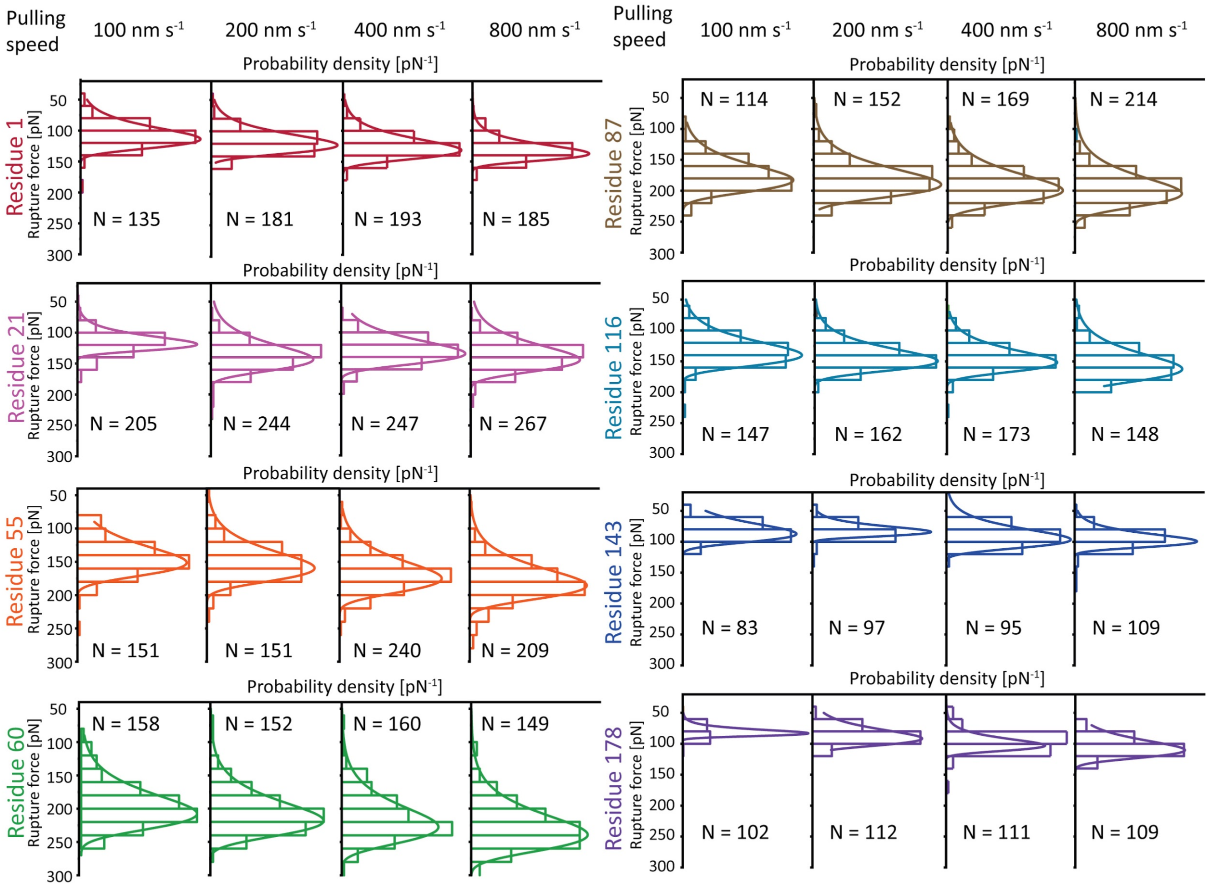


**Figure S5. Rupture force histograms of the anticalin:CTLA-4 complex under different pulling geometries.** The histograms were fitted with the closed form expression of the Bell-Evans model to extract the most probable rupture force.

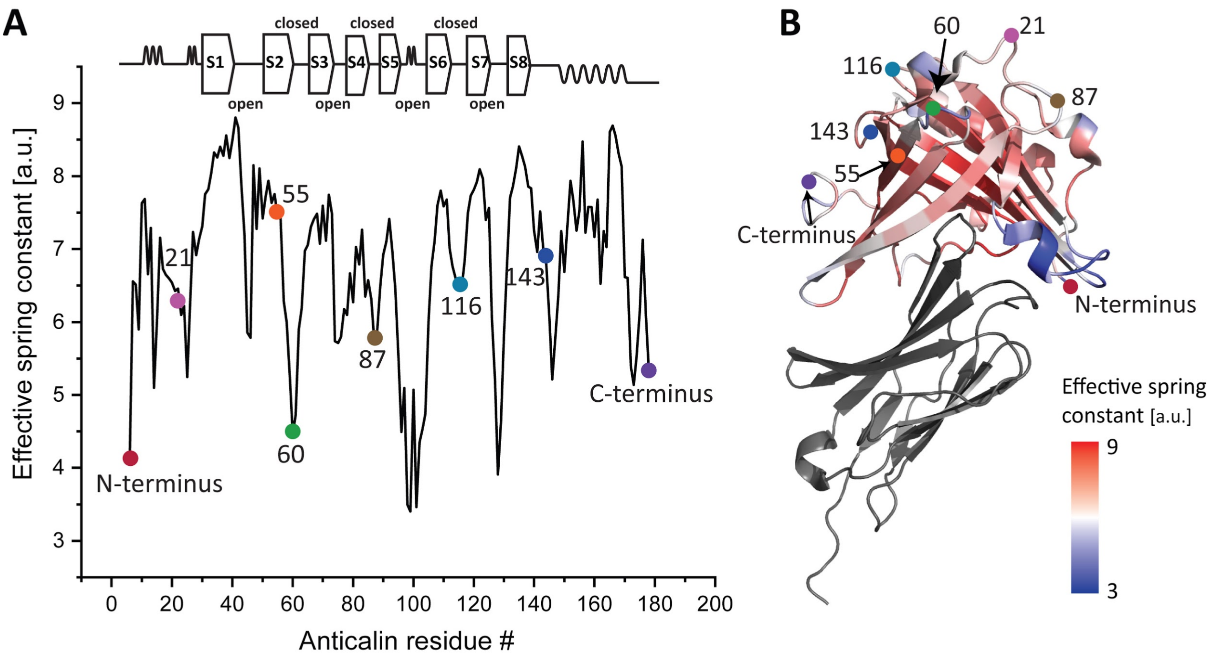


**Figure S6. Effective force constant between the C-terminus of CTLA-4 and all anticalin residues. A:** The effective force constant is plotted against the anticalin residue number and mapped to the secondary structure. Rigid regions (β-strands and helices) have higher force constants and the flexible regions have lower force constants. **B:** Heat map of the CTLA-4:anticalin structure (PDB 3BX7) showing the effective force constant between anticalin residues and the CTLA-4 C-terminus. The eight anchor residues on anticalin are highlighted.

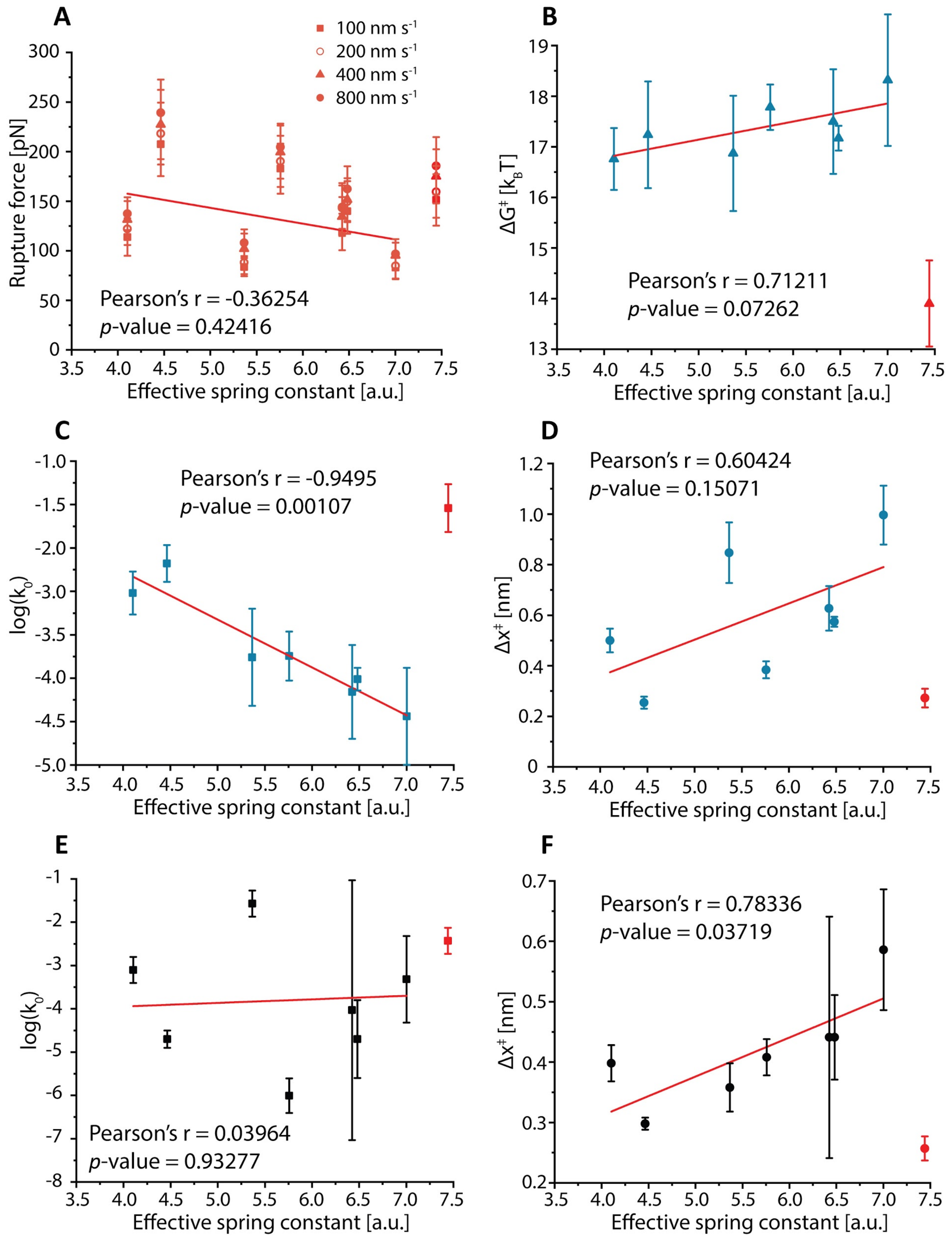


**Figure S7. Correlations between rupture force or unbinding energy profile and effective spring constant between anchor points. A:** the rupture force does not significantly correlate with the effective spring constant. **B-D:** the $\Delta G^{\ddagger}$ (**B**) and $\Delta x^{\ddagger}$ (**D**) calculated using DHS model are positively correlated with the effective spring constant while the $k_0$ (**C**) is negatively correlated with the effective spring constant. **E and F:** the $k_0$ calculated using BE model (**E**) does not have significant correlation with effective spring constant while the $\Delta x^{\ddagger}$ (**F**) is positively correlated.

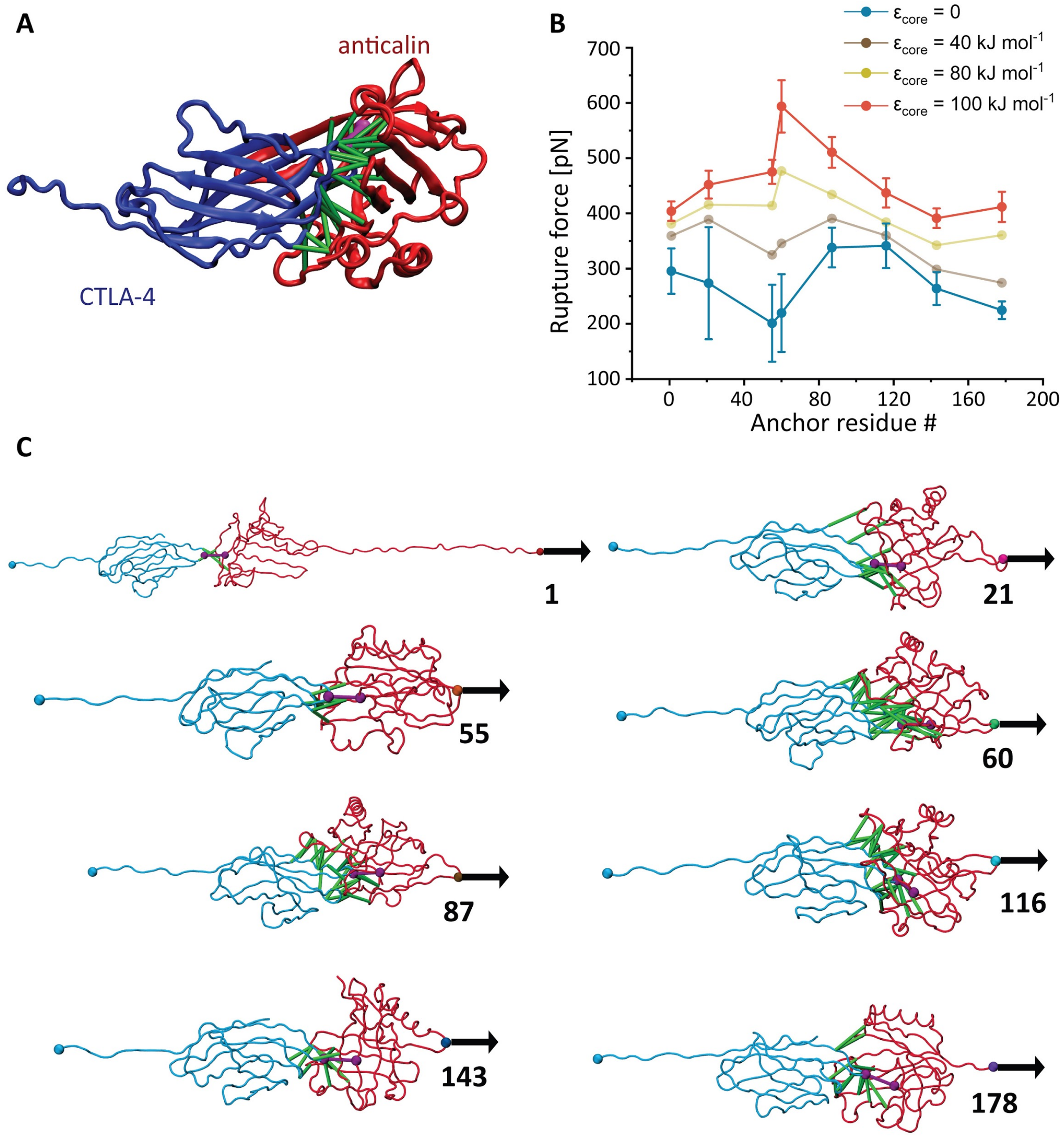


**Figure S8. Molecular dynamics (MD) simulations and *in silico* force spectroscopy. A:** Crystallographic structure of the anticalin:CTLA-4 complex (PDB code 3BX7) with CTLA-4 shown in blue and anticalin in red. High frequency native contacts ($\varepsilon_{Gō\text{-}MARTINI}$ = 9.414 kJ mol$^{-1}$ and $\varepsilon_{core}$ = 100 kJ mol$^{-1}$) determined in all-atom MD simulation that describe the protein-protein binding interface are shown as green and purple solid lines. **B:** Force-residue profile with $\varepsilon_{core}$ equal to 0.0 (blue), 40.0 (brown), 80.0 (yellow) and 100.0 (red) kJ mol$^{-1}$ with standard deviation as the error bars shown for $\varepsilon_{core}$= 0 and $\varepsilon_{core}$= 100. **C:** Different conformations of the complex are captured by coarse-grained simulations at the rupture force. The most relevant native contacts with $\varepsilon_{Gō\text{-}MARTINI}$ and $\varepsilon_{core}$ that contribute to the stability of the pulling geometry at the interface of the complex are highlighted as green and purple solid lines respectively. Number below the arrow indicates the anchor residue number.

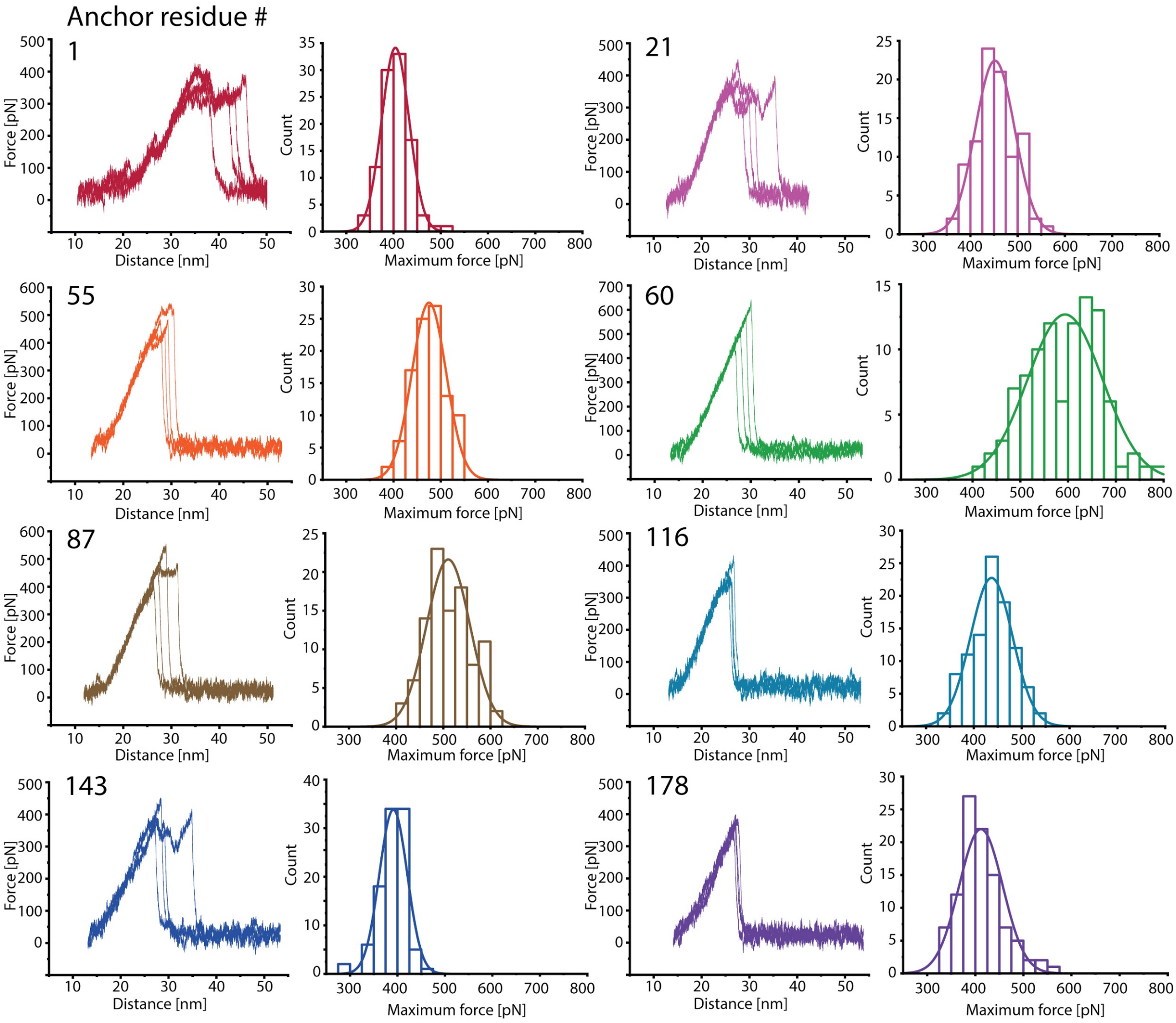


**Figure S9. Example force-extension curves and rupture force histograms of anticalin:CTLA-4 complex at different pulling geometries in Gō-MARTINI simulation.** The pulling studies were carried out at $5\times10^7$ nm $s^{-1}$. Each histogram was obtained based on an ensemble of 100 trajectories, with four of the trajectories shown here as examples.

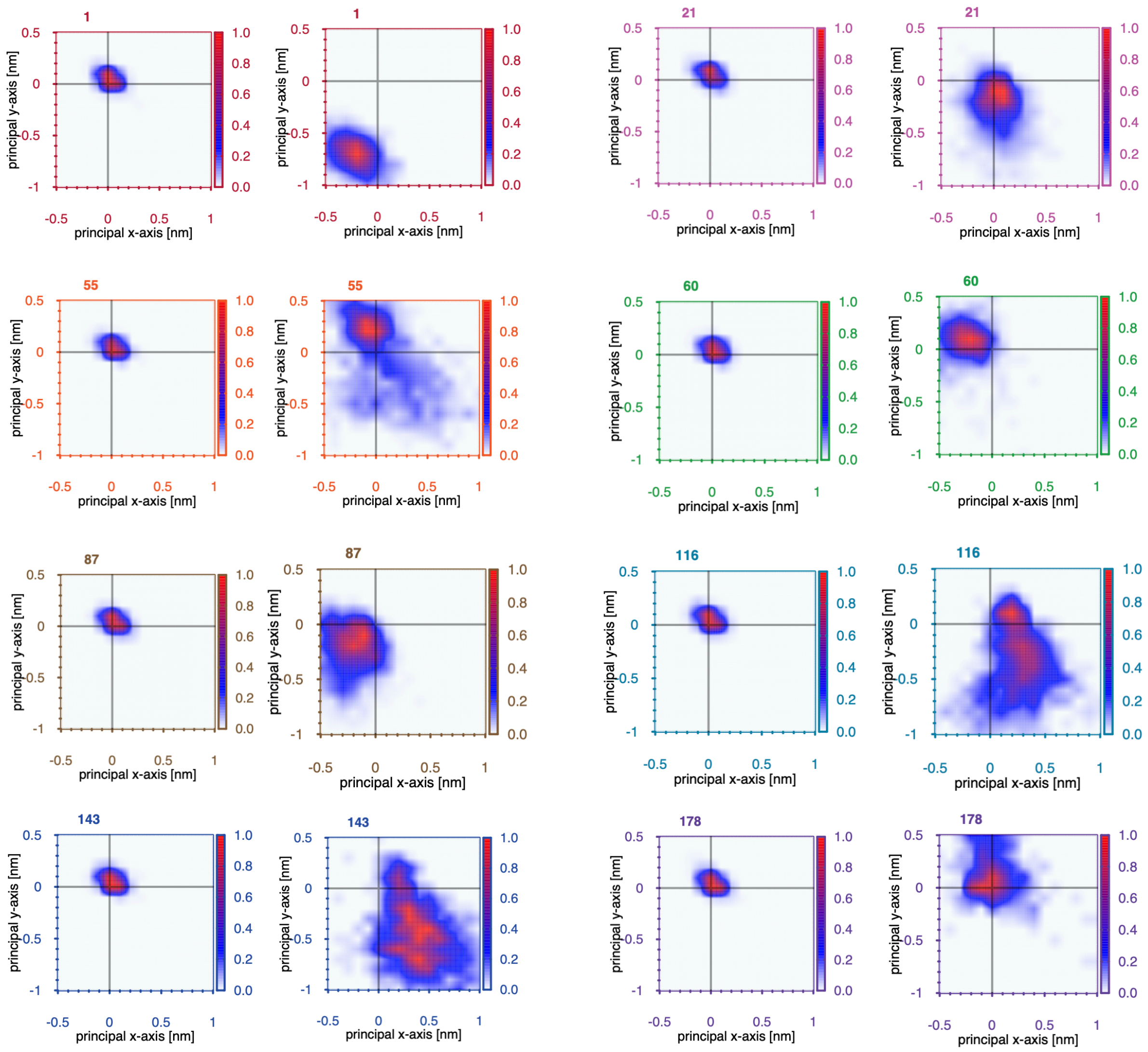


**Figure S10. Relative motion of the anticalin COM during Gō-MARTINI stretching simulations of the anticalin:CTLA-4 complex.** Data is presented for each pulling geometry at zero applied force (left side) and at $F_{max}$ (right side). Color bars indicate the probability to find the COM at a given position on the X-Y plane which is perpendicular to the z direction of symmetry of the complex. Numbers at the top left corner represent the anchor residue number.

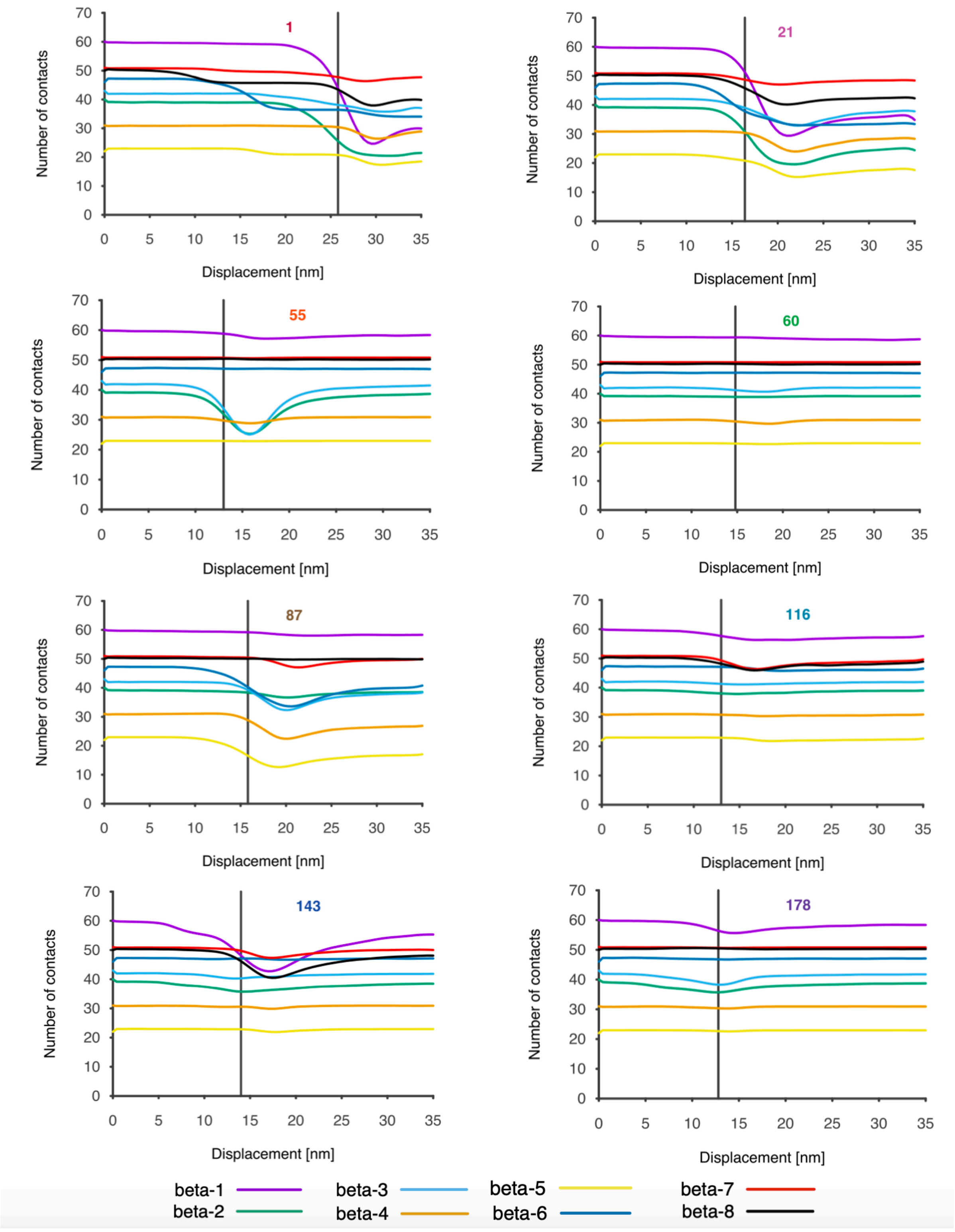


**Figure S11. Evolution of the anticalin intrachain native contacts (NC) during Gō-MARTINI stretching simulations of anticalin:CTLA-4 complex.** Each color line represents the set of NC for each beta-sheet of the anticalin. Large deviations from the original set involve partial unfolding and a gain in flexibility of the anticalin. Vertical black lines show the position of breaking of the interface contacts at rupture force. Numbers at the top represent the anchor residue number.

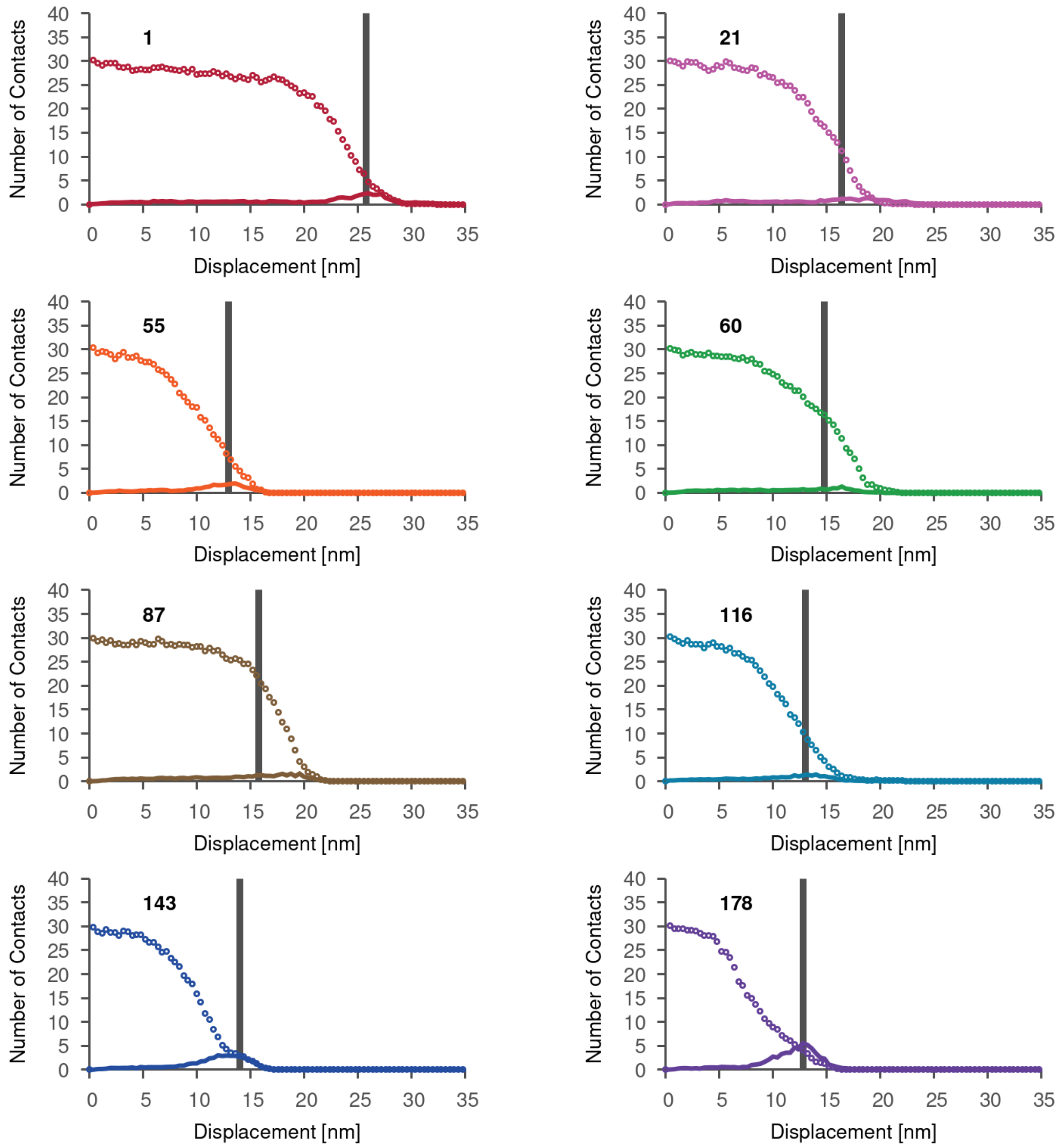


**Figure S12. Profile of native (open circles) and non-native (solid line) interface contacts during Gō-MARTINI stretching simulations of anticalin:CTLA-4 complex.** The vertical black line shows the position of the native contacts at rupture force. Native contacts and non-native contacts are calculated in the ensemble of pulling trajectories (n=100). Error bars are given by the size of the symbol. Number next to native contact profile represents the anchor residue number.

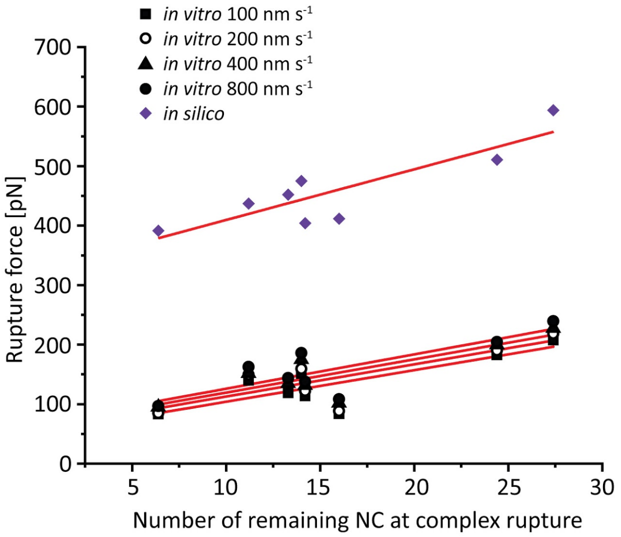


| Correlation | Pearson's r | *p*-value |
|---|---|---|
| *in vitro* 100 nm $s^{-1}$ | 0.818 | 0.013 |
| *in vitro* 200 nm $s^{-1}$ | 0.803 | 0.016 |
| *in vitro* 400 nm $s^{-1}$ | 0.831 | 0.011 |
| *in vitro* 800 nm $s^{-1}$ | 0.819 | 0.013 |
| *in silico* | 0.872 | 0.005 |

**Figure S13. Rupture forces *vs.* number of remaining NC at complex rupture.** The rupture forces measured both *in vitro* and *in silico* are positively correlated ($p<0.05$) with the number of remaining NC at complex rupture.

**Table S1** Effective spring constant between the C terminus of CTLA-4 and different anchor points on anticalin.

| Anchor residue # | 1 | 21 | 55 | 60 | 87 | 116 | 143 | 178 |
|---|---|---|---|---|---|---|---|---|
| Effective spring constant [a.u] | 4.10 | 6.43 | 7.44 | 4.46 | 5.76 | 6.48 | 7.00 | 5.37 |

**Table S2** Non-bonded (VdW and coulomb interactions) energy contribution for each protein chain in the anticalin:CTLA-4 complex and protein-protein interface at different levels of representation. Energy value is given in kJ/mol and next to it in parenthesis as a percentage of the total energy nonbonded energy in the system.

| Level of description | **CTLA-4** | **Interface** | **Anticalin** |
|---|---|---|---|
| All-atom MD | -2672.90 (37%) | -259.60 (4%) | -4255.20 (59%) |
| Gō-MARTINI | -3996.00 (36%) | -629.10 (6%) | -6410.70 (58%) |

**Table S3** Statistics of the total number of native contacts (NC) present in each protein component of anticalin:CTLA-4 complex at different pulling geometries. Average values of NC and the standard deviations are calculated in the ensemble of 100 pulling simulations at the rupture force.

| Anchor point on anticalin | **CTLA-4** | **Interface** | **Anticalin** |
|---|---|---|---|
| 1 | 185.0±2.2 | 14.2±11.5 | 200.2±30.1 |
| 21 | 185.0±2.3 | 13.3±8.5 | 228.0±25.3 |
| 55 | 184.0±3.1 | 14.0±7.2 | 240.2±13.0 |
| 60 | 179.1±6.0 | 27.4±3.0 | 265.1±3.4 |
| 87 | 181.0±5.0 | 24.4±4.4 | 241.0±18.0 |
| 116 | 184.3±2.4 | 11.2±7.0 | 250.1±10.0 |
| 143 | 184.3±2.4 | 6.4±5.0 | 231.0±22.0 |
| 178 | 185.0±2.1 | 16.0±4.0 | 252.0±5.0 |

**Table S4** Dissociation constants between CTLA-4 and anticalin mutants

| Anticalin mutant | I55AzF | E60AzF | E143AzF |
|---|---|---|---|
| Dissociation constant [nM] | 140±68 | 82±50 | 59±31 |

**Supplementary notes**

**Supplementary Note 1 Amino acid sequences and Addgene accession codes**

Color code: anticalin, CTLA-4, FLN, ELP, His-tag, ybbr tag, Fgβ, Streptag, SdrG

**CTLA4-FLN-ELP-His-ybbr (Addgene accession code 168038)**

HVAQPAVVLASSRGIASFVCEYASPGKATEVRVTVLRQADSQVTEVCAATYMMGNELTFLDDSICTGTSSGNQVNLTIQGLRAMDTGLYICKVELMYPPPYYLGIGNGTQIYVIDPEPGSGSGSGSADPEKSYAEGPGLDGGESFQPSKFKIHAVDPDGVHRTDGGDGFVVTIEGPAPVDPVMVDNGDGTYDVEFEPKEAGDYVINLTLDGDNVNGFPKTVTVKPAPGSGSGSHGVGVPGMGVPGVGVPGVGVPGVGVPGVGVPGVGVPGVGVPGVGVPGEGVPGEGVPGVGVPGMGVPGVGVPGVGVPGVGVPGVGVPGVGVPGVGVPGVGVPGEGVPGEGVPGVGVPGMGVPGVGVPGVGVPGVGVPGVGVPGVGVPGVGVPGVGVPGVGVPGEGVPGEGVPGWRGHHHHHHGSDSLEFIASKLA

**Anticalin-FLN-ELP-His-ybbr (Addgene accession code 168039)**

QDSTSDLIPAPPLSKVPLQQNFQDNQFHGKWYVVGLAGNRILRDDQHPMNMYATIYELKEDKSYNVTSVISSHKKCEYTIATFVPGSQPGEFTLGNIKSYGDKTSYLVRVVSTDYNQYAVVFFKLAEDNAEFFAITIYGRTKELASELKENFIRFSKSLGLPENHIVFPVPIDQCIDGGSGSGSGSADPEKSYAEGPGLDGGESFQPSKFKIHAVDPDGVHRTDGGDGFVVTIEGPAPVDPVMVDNGDGTYDVEFEPKEAGDYVINLTLDGDNVNGFPKTVTVKPAPGSGSGSHGVGVPGMGVPGVGVPGVGVPGVGVPGVGVPGVGVPGVGVPGVGVPGVGVPGEGVPGEGVPGVGVPGMGVPGVGVPGVGVPGVGVPGVGVPGVGVPGVGVPGVGVPGEGVPGEGVPGVGVPGMGVPGVGVPGVGVPGVGVPGVGVPGVGVPGVGVPGVGVPGVGVPGEGVPGEGVPGWRGHHHHHHGSDSLEFIASKLA

**Fgβ-anticalin-His-ybbr (Addgene accession code 168040)**

FFSARGHRPLDGSGSQDSTSDLIPAPPLSKVPLQQNFQDNQFHGKWYVVGLAGNRILRDDQHPMNMYATIYELKEDKSYNVTSVISSHKKCEYTIATFVPGSQPGEFTLGNIKSYGDKTSYLVRVVSTDYNQYAVVFFKLAEDNAEFFAITIYGRTKELASELKENFIRFSKSLGLPENHIVFPVPIDQCIDGGSGSGSHHHHHHGSDSLEFIASKLA

**Fgβ-StrepTag-DBCO peptide**

NEEGFFSARGHRPLDGSWSHPQFEKGSGSC-DBCO

**Anticalin mutant-His (Addgene accession codes 168041-168046)**

21 55 60

QDSTSDLIPAPPLSKVPLQQNFQDNQFHGKWYVVGRAGNTGLREDKDPGKMFATIYELKED
KSY

87 116

NVTSVISSHKKCEYTIATFVPGSQPGEFTLGNIKSYGDKTSYLVRVVSTDYNQYAVVFFKLAE
DNA

143

EFFAITIYGRTKELASELKENFIRFSKSLGLPENHIVFPVPIDQCIDGGSGSGSHHHHHH

Mutants for internal anchor point measurements (residues 21, 55, 60, 87, 116 or 143) were made by replacing the amino acid at the corresponding position with p-azido-L-phenylalanine (AzF) using amber suppression (see Methods-amber suppression section).

**SdrG-FLN-ELP-His-ybbr (Addgene accession code 168047)**

EQGSNVNHLIKVTDQSITEGYDDSDGIIKAHDAENLIYDVTFEVDDKVKSGDTMTVNIDKNT
VPSDLTDSFAIPKIKDNSGEIIATGTYDNTNKQITYTFTDYVDKYENIKAHLKLTSYIDKSKVP
NNNTKLDVEYKTALSSVNKTITVEYQKPNENRTANLQSMFTNIDTKNHTVEQTIYINPLRYSA
KETNVNISGNGDEGSTIIDDSTIIKVYKVGDNQNLPDSNRIYDYSEYEDVTNDDYAQLGNNND
VNINFGNIDSPYIIKVISKYDPNKDDYTTIQQTVTMQTTINEYTGEFRTASYDNTIAFSTSSGQG
QGDLPPEGSGSGSGSADPEKSYAEGPGLDGGESFQPSKFKIHAVDPDGVHRTDGGDGFVVTIE
GPAPVDPVMVDNGDGTYDVEFEPKEAGDYVINLTLDGDNVNGFPKTVTVKPAPGSGSGSHG
VGVPGMGVPGVGVPGVGVPGVGVPGVGVPGVGVPGVGVPGVGVPGEGVPGEGVPGVGVP
GMGVPGVGVPGVGVPGVGVPGVGVPGVGVPGVGVPGVGVPGEGVPGEGVPGVGVPGMGV
PGVGVPGVGVPGVGVPGVGVPGVGVPGVGVPGVGVPGEGVPGEGVPGWRGHHHHHHGSD
SLEFIASKLA

**Supplementary Note 2 Python code used to calculate the effective force constants**

```
#Python version 3.8.2
#Code adapted from http://prody.csb.pitt.edu/tutorials/mech_stiff/sm.html
from prody import *
from pylab import *
ion()
anticalin, header = parsePDB('3bx7', header=True)
calphas = anticalin.ca
anm = ANM('CTLA4_anticalin ANM analysis')
anm.buildHessian(calphas, cutoff=13.0)  #cutoff in Å
anm.calcModes(n_modes='all')
```

```
stiffness = calcMechStiff(anm, calphas)
calcStiffnessRange(stiffness)
writeVMDstiffness(stiffness, anticalin, [122], [1,20], filename='Forceconstants')
```
